# Super-resolution Left Ventricular Flow and Pressure Mapping by Navier-Stokes-Informed Neural Networks

**DOI:** 10.1101/2024.04.12.589319

**Authors:** Bahetihazi Maidu, Pablo Martinez-Legazpi, Manuel Guerrero-Hurtado, Cathleen M. Nguyen, Alejandro Gonzalo, Andrew M. Kahn, Javier Bermejo, Oscar Flores, Juan C. del Alamo

## Abstract

Intraventricular vector flow mapping (VFM) is a growingly adopted echocardiographic modality that derives time-resolved two-dimensional flow maps in the left ventricle (LV) from color-Doppler sequences. Current VFM models rely on kinematic constraints arising from planar flow incompressibility. However, these models are not informed by crucial information about flow physics; most notably the pressure and shear forces within the fluid and the resulting accelerations. This limitation has rendered VFM unable to combine information from different time frames in an acquisition sequence or derive fluctuating pressure maps. In this study, we leveraged recent advances in artificial intelligence (AI) to develop AI-VFM, a vector flow mapping modality that uses physics-informed neural networks (PINNs) encoding mass conservation and momentum balance inside the LV, and no-slip boundary conditions at the LV endocardium. AI-VFM recovers the flow and pressure fields in the LV from standard echocardiographic scans. It performs phase unwrapping and recovers flow data in areas without input color-Doppler data. AI-VFM also recovers complete flow maps at time points without color-Doppler input data, producing super-resolution flow maps. We show that informing the PINNs with momentum balance is essential to achieving temporal super-resolution and significantly increases the accuracy of AI-VFM compared to informing the PINNs only with mass conservation. AI-VFM is solely informed by each patient’s flow physics; it does not utilize explicit smoothness constraints or incorporate data from other patients or flow models. AI-VFM takes 15 minutes to run in off-the-shelf graphics processing units and its underlying PINN framework could be extended to map other flow-associated metrics like blood residence time or the concentration of coagulation species.

## 1 Introduction

Echocardiography is the most used imaging technique in the clinical setting to measure intracardiac flow, as it is non-invasive, portable, and widely available [1]. Modalities such as color-, pulsed-, and continuous-wave Doppler facilitate the bedside assessment of intracardiac blood flow by providing time-resolved velocities in the direction of the ultrasound beam. However, despite its well-known accuracy and accessibility, Doppler velocity measurements are unidirectional, often missing critical information about flow patterns.

Two- and three-directional vectorial maps of intracardiac flow can be acquired using phase-contrast cardiac magnetic resonance imaging (PC-MRI) [2–5]. However, compared with echocardiography, PC-MRI has limited temporal and spatial resolutions, involves long acquisition times, and is less accessible, which can hamper its application in clinical practice [6–8]. Computed tomography (CT) is more widely available, can assess cardiovascular anatomy with high spatial resolution in the imaging plane, and offers the possibility to obtain volumetric data. However, no analysis tools quantify flow directly from CT images. While computational fluid dynamics (CFD) analysis can be performed using CT-derived segmentations [9], CFD is critically sensitive to modeling choices and boundary conditions.

Given its potential value in the diagnosis and prognosis of heart failure, ultrasound flow mapping inside the left ventricle (LV) has received significant attention. Several ultrasound methods have been developed to capture LV flow, balancing temporal and spatial resolutions with clinical accessibility. Echo particle image velocimetry (Echo-PIV) can obtain time-resolved 2D intraventricular velocity fields with good spatial and temporal resolution from contrast-infused images [10, 11], and 3D extensions have been proposed [12]. However, its sensitivity to operational parameters like image quality, frame rate, insonation angle, and absolute flow velocity [13], and the consequent need for a finely tuned contrast infusion, have hindered the clinical application of Echo-PIV. Emerging modalities like Blood Speckle Imaging (BSI) use analyses analogous to PIV to the information in the high-frequency ultrasound scattered by red blood cells, simplifying LV flow mapping. However, this technique has been limited so far by the shallow penetration of high-frequency ultrasound. Although its feasibility has been demonstrated in adults [14], BSI is most often applied in the pediatric setting [15].

In the past decade, vector flow mapping (VFM) has gained acceptance as a modality to obtain time-resolved 2D velocity maps from color-Doppler echocardiographic acquisitions in the LV [16–22], as well as 3D velocity maps from triplane color-Doppler [23]. Early implementations of VFM recovered the cross-beam velocity component by enforcing planar mass conservation along circular arcs of the color-Doppler sector [16–18]. Subsequent extensions reformulated VFM as a maximum-likelihood estimation problem with priors based on mass conservation and noise reduction [19, 20]. Other VFM variants use the stream function-vorticity formulation [22, 24]. Despite these advances, existing VFM methods do not incorporate physical information about momentum balance (i.e., the Navier-Stokes equations) and remain sensitive to imaging artifacts that lead to spatial and temporal gaps in the color-Doppler input images. Moreover, current VFM implementations focus exclusively on recovering the cross-beam velocity, overlooking significant flow variables like pressure, which has to be recovered by secondary analyses [25]. The dynamical interplay between pressure and flow acceleration is not used to inform current VFM methods.

In parallel with these efforts, artificial intelligence (AI) models based on deep learning (DL) have swiftly transformed cardiovascular image analysis [26]. Initially focused on automating the interpretation of echocardiograms and electrocardiograms or the segmentation of cardiovascular structures [27–29], these models can enhance flow imaging data by removing phase-wrapping artifacts, reconstructing complete fields from scattered measurements, and increasing the resolution of flow measurements [30–32]. Physics-informed neural networks (PINNs) harness recent advancements in DL to infer hidden information in flow measurements constrained by the governing equations of fluid mechanics [33, 34]. A notable feature of PINNs is that they can be trained on sparse data and in a patient-specific manner, making them well-suited for VFM. The application of DL models or, more specifically, PINNs to VFM remains underexplored despite noteworthy ongoing work [35].

This manuscript introduces AI-VFM, a vector flow mapping method that uses PINNs to reconstruct the cross-beam flow velocity and fluctuating pressure in the apical long-axis view of the left ventricle. Like other VFM implementations, AI-VFM uses a color-Doppler sequence and a time-resolved delineation of the LV endocardial wall as inputs. Its outputs are a phase-unwrapped color-Doppler map together with reconstructed cross-beam velocity and fluctuating pressure maps. The PINN is trained for each patient-specific acquisition to minimize a loss function that aggregates the difference between the input and output Doppler maps, as well as the residuals of the continuity and Navier-Stokes equations in the imaging plane, including boundary conditions at the endocardial border. Most notably, AI-VFM leverages momentum balance to recover hidden information about flow dynamics which was inaccessible to previous methods.

The manuscript is organized as follows. Section 2 describes the AI-VFM implementation and the methodology for validation and sensitivity analyses using ground-truth data from CFD simulations. Section 3 reports results from those analyses and illustrates the application of AI-VFM to echocardiographic acquisitions, including cases with evident missing information (“holes”) or artifacts in the input Doppler images. Section 4 discusses the strengths and weaknesses of AI-VFM, suggests applications of this modality, and outlines the potential extensions of AI-VFM.

## 2 Materials and Methods

### 2.1 Governing Equations of LV flow

This work relies on mathematical models of LV flow to define loss function terms in the physics-informed neural network (PINN) and to generate synthetic ground-truth data for validation via computational fluid dynamics (CFD) analysis. These models use the same governing equations of fluid flow but differ in some aspects, like their physical domains and coordinate systems (see §2.2 and §2.6 below). In all cases, blood was modeled as an incompressible, Newtonian fluid of constant kinematic viscosity ν = 3 *×* 10*^−^*^6^, m^2^/s. Consequently, the velocity ^(⃗^**v**) and pressure (p) fields were assumed to obey the continuity and Navier-Stokes equations, i.e.,

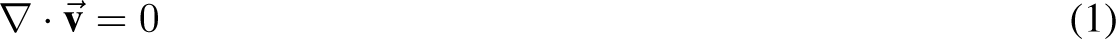

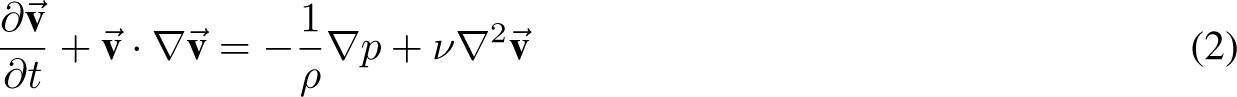

where ρ is the density of blood. The boundary conditions for these equations are no-slip at the endocardial wall, i.e., ⃗**v** = ⃗**v***_wall_*.

### 2.2 LV geometrical domain for AI-VFM

In AI-VFM, we adopted a planar polar coordinate system with its origin at the ultrasound transducer (Figure 1A), as customary in the VFM literature [18]. In this reference frame, the radial velocity component v*_r_* is parallel to the ultrasound beam and, thus, measurable by color-Doppler. In contrast, the azimuthal velocity component, v*_θ_*, and the pressure field p are unknown. The equations of fluid motion (1–2) were enforced on a domain Ω corresponding to an LV segmentation obtained from echocardiographic imaging as described in §2.9 below. Boundary conditions were imposed on the endocardial border of the segmentation, ∂Ω.

**Figure 1:**
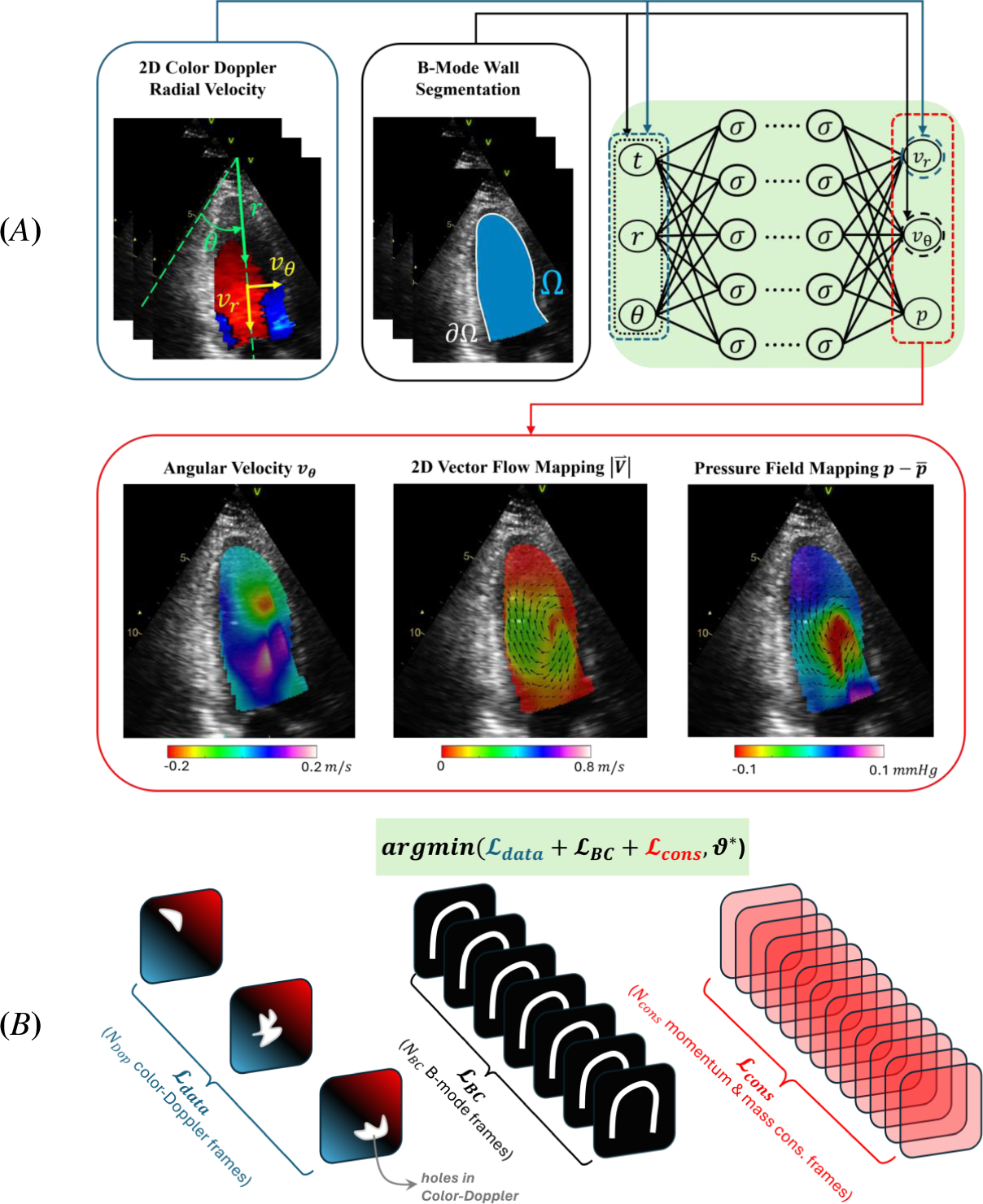
AI-VFM flowchart. Raw color Doppler radial velocity field is used for training. B-mode wall segmentation to obtain LV boundary points to impose the boundary condition. A fully connected neural network is employed to infer radial velocity *v_r_*, angular velocity *v_θ_*, and pressure field *p*. The neural network is penalized by a loss function constructed from training data *L_data_* and underlying physical laws *L_BC_* and *L_cons_*. The total loss *L_total_* is minimized using the *Adam* optimizer. Neural network parameters (e.g., weights *ϑ^∗^*) are updated via backpropagation during each iteration.

### 2.3 Color-Doppler Phase Unwrapping in AI-VFM

During Doppler acquisitions, phase wrapping (i.e., aliasing) artifacts occur when blood velocity exceeds the maximum detectable velocity (e.g., Nyquist velocity), causing the detected Doppler phase to wrap around 2π and altering the sign and magnitude of the measured velocity, i.e.,

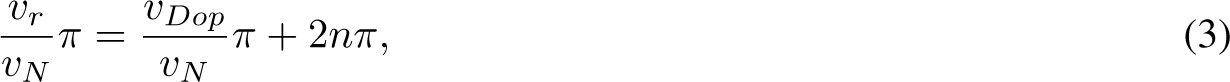

where v*_r_* is the actual velocity in the direction parallel to the ultrasound beam, v*_Dop_* is the measured Doppler velocity, v*_N_* is the Nyquist velocity, and n is the number of phase wraps. Since aliased velocities do not obey the equations of fluid motion (1-2), Doppler measurements cannot be used directly as training data in AI-VFM. However, trigonometric functions (e.g., sines and cosines) of both sides of equation 3 are equal. Thus, we used sin(πv*_r_*/v*_N_*) and cos(πv*_r_*/v*_N_*) as training data fields in our neural network. A similar approach was previously implemented in PINN to perform phase-unwrapping in 4D flow MRI velocities [36]. We found this method yielded flow and pressure fields free of aliasing artifacts with no need for additional constraints and without considering n into the PINN.

### 2.4 Neural Network Architecture and Super-Resolution Loss Function

AI-VFM (Figure 1A) uses physics-informed neural networks (PINNs) to reconstruct the cross-beam flow velocity and the fluctuating pressure from a color-Doppler sequence and a time-resolved delineation of the LV endocardial wall. A PINN is a neural network trained to minimize a residual including physical laws. Therefore, it learns the training data and the underlying (i.e., hidden) dynamics of the system. In AI-VFM, we use a fully connected neural network to approximate the following mapping function: [*t*, *r*, θ] *1→* [v*_r_*, v*_θ_*, *p*], where the inputs are time and the spatial coordinates of the points inside the LV, and the outputs are the corresponding velocities (radial and cross-beam components) and pressure fluctuations. Input and output variables are defined on N*_out_* frames along the cardiac cycle. In all the cases we studied, we adopted a neural network configuration with 10 layers, each consisting of 150 neurons. This configuration aimed to balance low expressivity (shallow layer) and overfitting (deep layer). Given that the underlying physical constraints involve nonlinear equations, we employed nonlinear *swish* activation functions [37]. The neural network weights were initialized using Xavier’s scheme [38], and weight normalization was employed to accelerate training [39].

The loss function employed to train the AI-VFM PINN comprises three terms representing the differences with respect to the training data (*L_data_*), the wall boundary conditions (*L_BC_*), and the governing mass and momentum conservation equations (*L_cons_*),

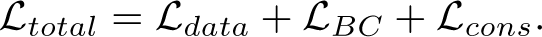

Color-Doppler v*_r_* fields usually have different temporal resolutions than the B-mode acquisitions used to delineate the LV wall, where boundary conditions are imposed. To account for this disparity, and to achieve temporal super-resolution in the output flow and pressure fields, we devised the strategy outlined in Figure 1B. The data loss, *L_data_*, was sampled in the available N*_Dop_* color-Doppler frames, whereas the boundary condition loss, *L_BC_*, was sampled in the available N*_BC_* B-mode frames, and the governing equations loss, *L_cons_*, was sampled at the N*_cons_* frames. We then set the number of output temporal frames, N*_out_*, equal to N*_cons_* > N*_BC_* > N*_Dop_* to achieve temporal super-resolution. The data loss term was defined as

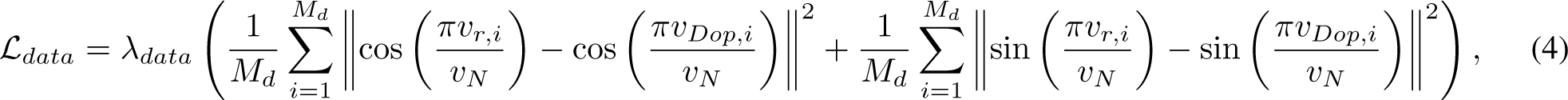

where λ*_data_* is a hyperparameter and i indicates members within a list of M*_d_* sampled points in the segmented LV mask (Ω) across the available N*_Dop_* color-Doppler frames. We accounted for possible holes in the color-Doppler measurements caused by, e.g., low signal power or imaging artifacts, by enabling the selective removal of subdomains of Ω in the calculation of *L_data_*. Therefore, the sampled points used to compute *L_data_* were randomly redrawn each training iteration, excluding points within data holes. The boundary condition loss, weighed by the hyperparameter λ*_BC_*, included the residual of the no-slip boundary condition sampled at M*_bc_* points of the LV boundary (∂Ω) across the available N*_BC_* bright-mode frames,

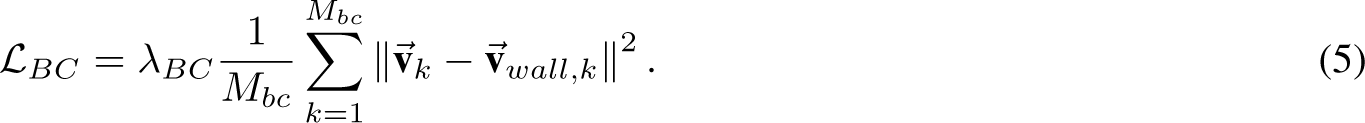

Finally, the mass and momentum conservation loss was defined as

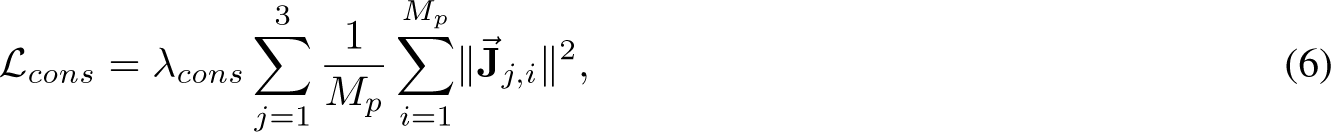

where the vector **J⃗***_j,i_* = J*_j_*(r*_i_*, θ*_i_*, t*_i_*) is formed by the residuals of the continuity equation and the radial and azimuthal components of the non-dimensional Navier-Stokes equations in polar coordinates, i.e.,

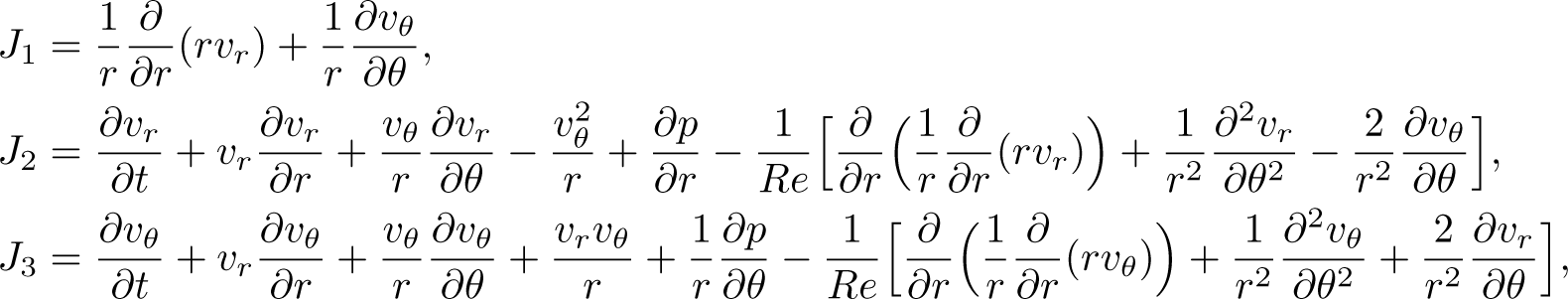

where Re = UD/ν is the Reynolds number obtained from the typical values of the diastolic jet velocity (U), the mitral valve diameter (D), and blood viscosity (ν). These loss terms were sampled at M*_p_* points randomly selected across N*_cons_* frames. The points were chosen inside the time-dependent Ω domain obtained by interpolating the LV mask segmentation from N*_BC_* frames to N*_cons_* frames. The temporal and spatial derivatives appearing in the definition of ^J⃗^*_j,i_* were computed using automatic differentiation [40] on the neural network.

To non-dimensionalize the flow variables used in training, the inputs (e.g., *t*, *r*) and training data (v*_r_*) were normalized by the characteristic scales specified in **Table 1**, and then back-dimensionalized during the inference stage after training is complete. Additionally, we employed a renormalization scheme to ensure all training variables had zero mean and unit variance. This step promotes training robustness and mitigates the issue of vanishing gradients during backpropagation [38].

**Table 1:** Characteristc scales used to non-dimensionalize inputs. Reynolds number defined by Re = UD/ν, where ν = 3 *×* 10*^−^*^6^m^2^/s is the kinematic viscosity, and U and D are the peak diastolic jet velocity and the diameter of the mitral valve, respectively. Spatial resolution for CFD benchmark and echocardiographic data. Case-specific N*_Dop_*, N*_BC_*, and N*_cons_* = 200. Iterations of AI-VFM for each example.

| Example | | $U$ [cm/s] | $D$ [cm] | Re | $\Delta r$ [mm] | $\Delta \theta$ [°] | $N_{Dop}$ | $N_{BC}$ | $N_{cons}$ | Iterations |
| --- | --- | --- | --- | --- | --- | --- | --- | --- | --- | --- |
| CFD |  | 50.00 | 2.73 | 4550 | 0.50 | 1.00 | 15 | 100 | 200 | 80K |
| ECHO | #1 | 73.60 | 3.57 | 8758 | 0.54 | 1.34 | 17 | 64 | 200 | 70K |
|  | #2 | 57.24 | 2.31 | 4407 | 0.54 | 1.34 | 19 | 74 | 200 | 68K |
|  | #3 | 57.24 | 2.40 | 4579 | 0.54 | 1.35 | 19 | 83 | 200 | 65K |

### 2.5 PINN runs

In all the PINN runs reported in this manuscript, a batch size of 10K sampling points (e.g., M*_d_* = M*_p_* = 10^4^) was used during each iteration to evaluate the losses with respect to training data and physical constraints, and the whole batch (i.e., all available points) was used to impose boundary conditions. The entire batch size for boundary conditions varied for CFD and ECHO depending on spatiotemporal resolution, with typical values being M*_bc_ ≈* 59K and 23K for CFD and ECHO, respectively. In our CFD validation, we chose λ*_data_* = 10, λ*_cons_* = 1, and λ*_BC_* = 10, whereas we set all the weights to be unity in the clinical application of Doppler data. The PINNs were run until *L_data_*, *L_BC_*, and *L_cons_* reached plateaus, as shown in Figure 2A-B. The number of iterations run for each case is indicated in **Table** 1. The total loss function was minimized using the

**Figure 2:**
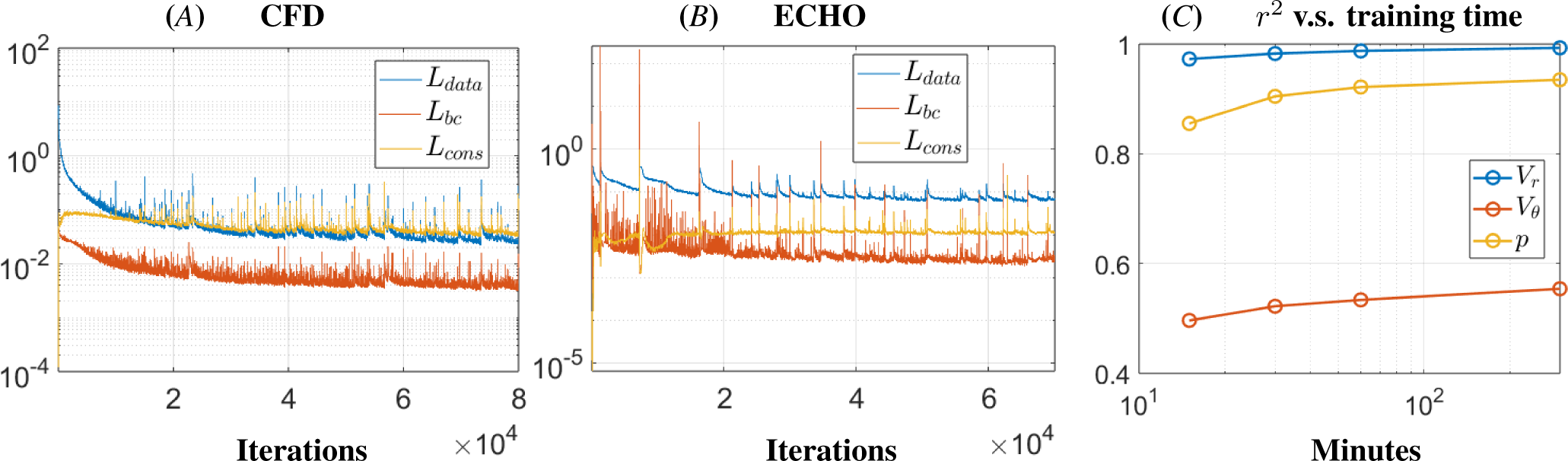
*A*) Loss values vs. AI-VFM training iteration using settings for CFD validation (Δ*r* = 0.5mm, Δ*θ* = 1.0°, *NDop* = 15, *NBC* = 100, and *Ncons* = 200); *B*) Loss values vs. training iterations using settings typical for clinical application of AI-VFM (Δ*r* = 0.54mm, Δ*θ* = 1.34°, *NDop* = 17, *NBC* = 64, and *Ncons* = 200); *C*) Accuracy vs. computational cost analysis in high-resolution CFD case (e.g., Δ*r* = 0.5mm, Δ*θ* = 0.3°) using *NDop* = *NBC* = *Ncons* = 200.

*Adam* adaptive optimization algorithm [41] with a default learning rate of 0.001. AI-VFM was implemented in Python using TensorFlow [42] library as described in Raissi et al. 2020 [43]. The training was performed using an NVIDIA A40 GPU with 48GB RAM, providing favorable accuracies in the recovered pressure and velocity fields within 15 mins, which improved accuracies an additional *≈* 10% by extending training runs to one hour (Figure 2C). All other pre-and postprocessing was done in MATLAB.

The classic VFM implementation relies on mass conservation but does not enforce momentum balance. To help understand how this modeling choice affects the accuracy of AI-VFM, we compared two AI-VFM implementations. In *kinematic* AI-VFM, we only included the continuity loss, J_1_, in the definition of *L_cons_*. In *dynamic* AI-VFM (or simply AI-VFM), we enforced momentum balance in addition to mass conservation as physical constraints, using the definition for *L_cons_* given in equation (6). Moreover, since pressure does not enter the continuity equation for an incompressible fluid, *kinematic* AI-VFM did not recover p.

### 2.6 LV geometrical domain for CFD

The LV cavity of our CFD simulation data was modeled as a half-ellipsoid (Figure 3A) with the endocardial surface given by

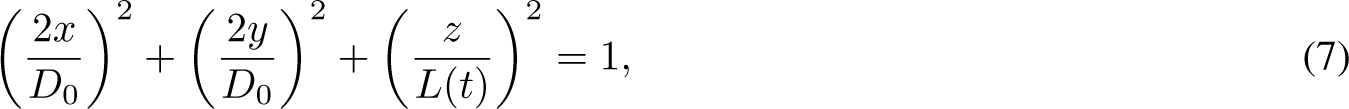

**Figure 3:**
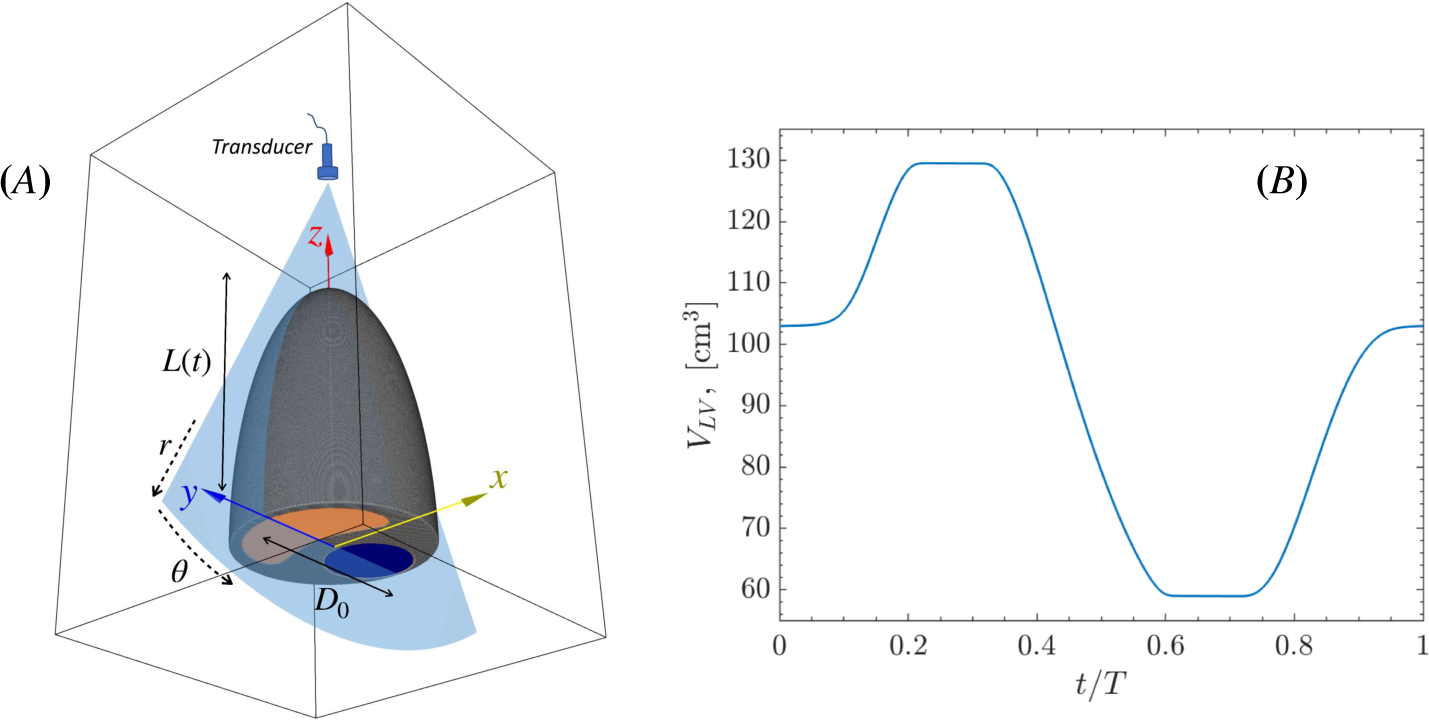
*A*) Sketch of the computational domain and idealized LV geometry, showing the mitral valve orifice in orange and the aortic valve orifice in blue; *B*) Time evolution of the LV volume prescribed in CFD simulations.

where D_0_ = 5.57 cm is the diameter of the LV base and L(t) is the time-dependent LV long-axis dimension (i.e., the distance between the LV base and its apex). LV beating in sinus rhythm at 70 b.p.m. was modeled by varying L(t) between L_min_ = 3.62 cm and L_max_ = 7.96 cm, yielding the LV volume vs. time evolution shown in Figure 3B. The two increments of LV volume correspond to the E and A waves mediated by LV relaxation and atrial contraction, with a physiological E/A wave ratio of 1.1. LV ejection fraction (EF = 0.54) and sphericity index (SpI = 0.49) were also set within physiological ranges [44, 45].

The openings at the LV base shown in Figure 3A represent the aortic and mitral orifices. They were modeled without leaflets and with two states: fully open or fully closed. The aortic orifice was modeled as a circle of diameter D*_AO_* = 2.52 cm centered at x*_AO_* = 0 and y*_AO_* = *−*1.48 cm (i.e., separated by a distance 0.05 cm to the edge of the LV base), yielding an area A*_AO_* = 4.97 cm^2^. A half-moon-shaped mitral orifice was defined by the boundary line:

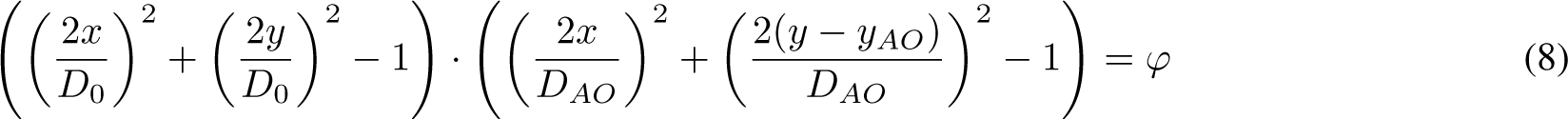

where the parameter φ was chosen to yield an orifice area A*_MO_* = 7.45 cm^2^.

### 2.7 CFD runs

Following previous works [46], the blood flow simulations of the idealized LV configuration described above were performed using TUCAN, an in-house GPU flow solver [47–50]. TUCAN uses a fractional step method to solve the Navier-Stokes equations for an incompressible fluid. The spatial discretization is based on second-order finite differences in a Cartesian staggered grid with constant grid spacing in all directions, Δx. Time integration is performed using a three-step, semi-implicit Runge-Kutta. The presence and motion of the LV walls are modeled in TUCAN using the immersed boundary method [51].

The simulations were initialized with zero velocity inside the LV and run for 5 cardiac cycles to ensure a quasi-periodic and converged flow. All the data presented in this paper correspond to the 5th cycle, for which 200 uniformly-spaced velocity and pressure fields were stored. For the present work, we chose Δt = 1.47 *×* 10*^−^*^4^ s and Δx = 0.5 mm, yielding a CFL < 0.6 throughout the cardiac cycle. These temporal and spatial resolution values agree with recent convergence studies for intracardiac flows [52].

### 2.8 Validation and Sensitivity Analysis on CFD-generated LV flow

Velocity and pressure fields from CFD simulations were interpolated into an 11 cm *×* 60 degree fan-shaped sector mimicking the long-axis apical echocardiographic view (Figure 3A). In this scheme, the ultrasound transducer would be located at the sector’s vertex, which also coincides with the origin of the polar coordinate system used in the VFM model (Figure 1A). To match the spatial resolution of standard color-Doppler acquisitions (see §2.9), the CFD data were interpolated using radial and azimuthal step sizes Δr = 0.5 mm and Δθ = 1.0*^◦^*, yielding a mesh with 220 *×* 50 points. Also, 200 temporal frames of flow fields evenly distributed over one cardiac cycle were obtained for training and validation.

To test the accuracies of *dynamic* and *kinematic* AI-VFMs’ when varying the temporal resolution of the input color-Doppler sequence, we first considered using all available training data (e.g., N*_Dop_* = N*_BC_* = N*_cons_* = 200). We then tested *dynamic* AI-VFM’s ability to provide super-resolution against more realistic temporal resolution of wide-sector color-Doppler and bright-mode acquisitions (e.g., N*_Dop_* = 15, N*_BC_* = 100, N*_cons_* = 200). To evaluate AI-VFM’s ability to recover hidden information in untrained frames, we ran another *dynamic* AI-VFM using N*_Dop_* = N*_BC_* = N*_cons_* = 15 and linearly interpolated the results to populate the velocity field up to 200 frames. The sensitivity of AI-VFM with respect to imaging resolution was tested by considering down-sampled cases with Δr = 1mm, Δθ = 1.3, 1.8*^◦^* and one up-sampled case Δr = 0.5mm, Δθ = 0.3. We also tested whether increasing N*_Dop_* from 15 to 50, 100, or 200 improved the accuracy of the recovered fields in both *dynamic* and *kinematic* models.

Aliasing artifacts were simulated by introducing a Nyquist limit of v*_N_* = 50 cm/s in v*_r_*. In addition, we investigated the effect of transducer misalignment with the LV and outflow jets by rotating the virtual Doppler sector as described in Figure 6(E & F). When varying one parameter, all other parameters are kept fixed at nominal values (i.e., Δr = 0.5mm, Δθ = 1.0°, N*_Dop_* = 200, and α = 0).

To compare the recovered flow fields vs. the ground-truth CFD data, we computed the coefficient of determination, r^2^, and the normalized root-mean-squared error (NRMSE) as follows:

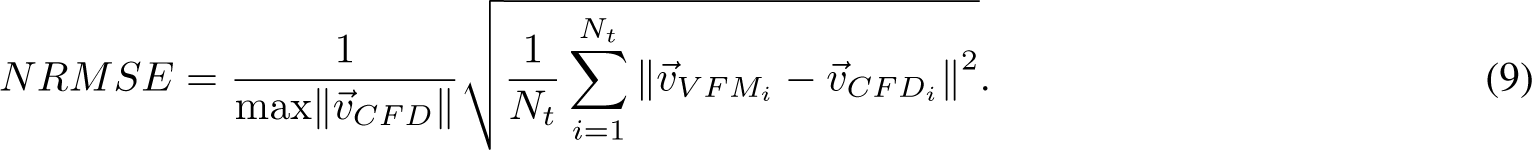

### 2.9 Echocardiographic Imaging

To assess the clinical feasiblity of AI-VFM, color-Dopper image sequences were acquired using a Vivid 7 scanner with broadband transducers (GE Healthcare) from three patients in sinus rhythm, one with dilated cardiomyopathy (patient #1) and two with acute myocardial infarction (patient #2 & #3). Standard B-mode and Doppler data were obtained as recommended in established guidelines [53]. Pulsed-wave Doppler spectrograms were obtained at the mitral valve tips and LV outflow tract, from which key cardiac cycle events were identified using EchoPac software (version 214, GE Healthcare) to temporally align data for postprocessing. Color-Doppler images covering the entire LV chamber were obtained in the apical three-chamber view (*∼* 10 cycles, frame rate *≈* 15 Hz), followed by the acquisition of 2D cine-loops (*∼* 3 cycles, frame rate > 60 Hz). The LV myocardial wall was segmented from the LV B-mode series using speckle-tracking software to delineate the endocardium (EchoPac, version 214, GE Healthcare). The spatial and temporal resolutions of the acquired Doppler images (e.g., v*_r_*) and flow parameters used in AI-VFM for training (N*_Dop_*), imposing boundary condition (N*_BC_*), and mass and momentum conservation equations (N*_cons_*) are summarized in **Table** 1.

### 2.10 Reference (Vanilla) VFM and Phase-Unwrapping Methods

For reference in our validation study, we also derived time-resolved 2D blood velocity fields from color-Doppler data inside the LV using Garcia *et al.*’s original method (as described in references [16,54], and hereon referred to as vanilla VFM). The vanilla VFM algorithm is fed by a color-Doppler acquisition and integrates the planar continuity equation, imposing no-penetration boundary conditions at the LV endocardium. Retrospective frame interleaving from multiple cycles and Fourier interpolation using 10 *−* 20 modes are applied to produce output sequences with 100-200 frames/beat. Data from heartbeats with > 5% heart rate variation are rejected. We used Loecher *et al.*’s one-step Laplacian correction [55] as the reference method for phase unwrapping of aliased color-Doppler values. In this method, a Laplacian equation is formulated for the number of phase wraps,

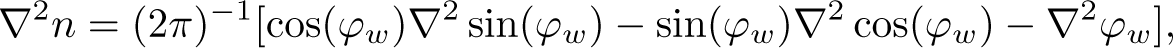

where φ*_w_* = πv*_w_*/v*_N_* is the wrapped phase and v*_w_* the aliased velocity. The solution to this equation is rounded off to the nearest integer number and plugged into equation 3 to correct the velocity field.

## 3 Results

This section presents validation and sensitivity analyses of AI-VFM using CFD-generated data. We also illustrate the clinical application of AI-VFM to improve ultrasound flow data. Overall, AI-VFM accurately recovers the cross-beam velocity component and fluctuating pressure inside the LV at temporal resolution higher than that of the training Doppler data. It also deals with aliasing artifacts and recovers missing velocity values.

### 3.1 AI-VFM Validation on CFD-generated LV flow

Figure 4 displays snapshots of LV flow velocity and fluctuating pressure fields during the early filling, late filling, and ejection phases of the cardiac cycle. This figure includes the CFD-generated ground-truth data and the results from two AI-VFM runs. In the first run, AI-VFM was trained using N*_cons_* = N*_BC_* = N*_Dop_* = 200 temporal frames, including the three timepoints represented in Figure 4. In the second run, AI-VFM was trained using a more realistic setting with N*_cons_* = 200, N*_BC_* = 100, N*_Dop_* = 15. Of note, in this second AI-VFM run, none of the color-Doppler data from the three time points represented in Figure 4 was used for training. Still, AI-VFM inferred the entire velocity field at these time points, thereby increasing the temporal resolution of the original data.

**Figure 4:**
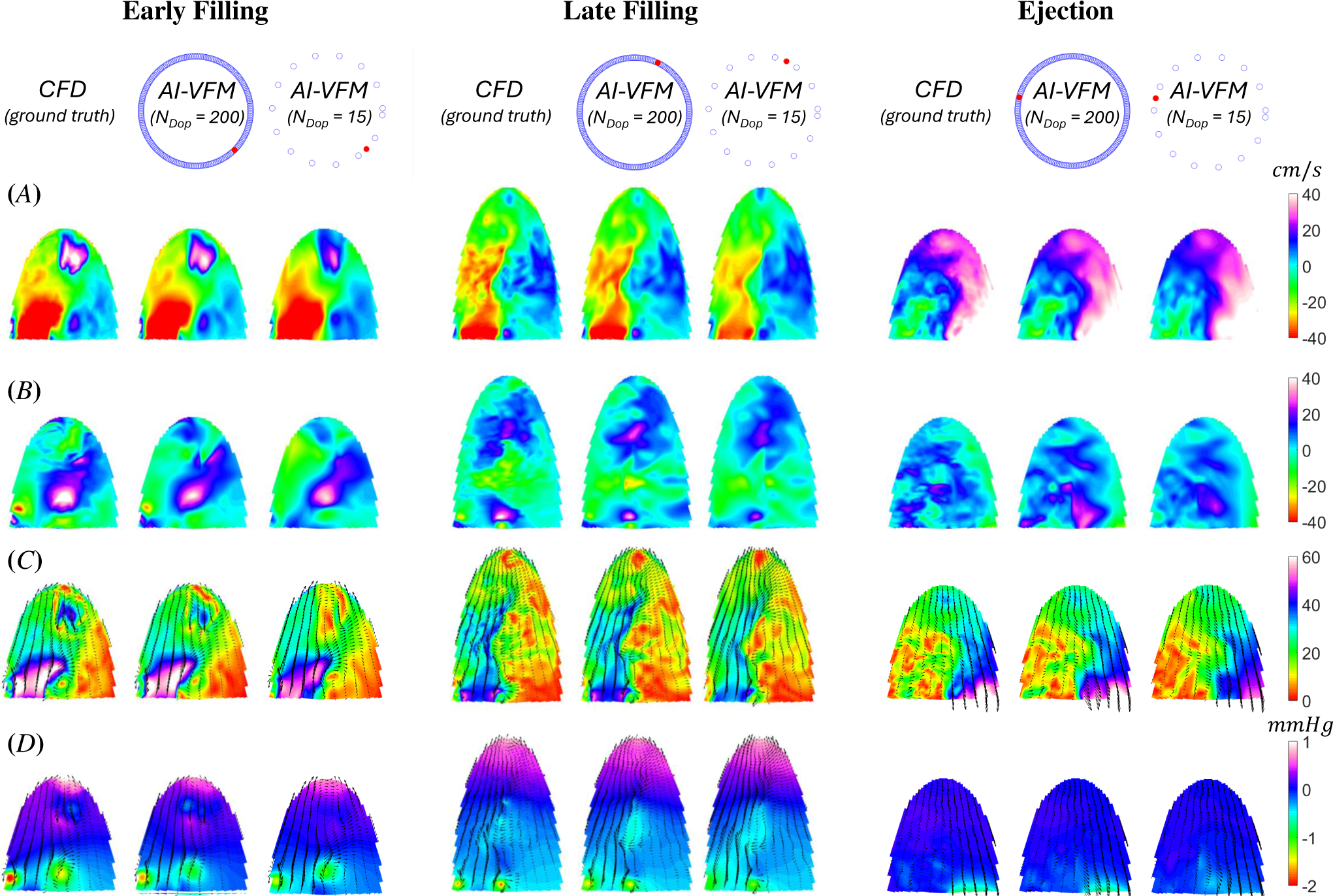
Velocity and pressure maps inferred by AI-VFM compared to ground-truth CFD data at early filling, late filling, and ejection (i.e., *t* = 0.85, 0.45, 0.21*T*, where *T* is the cardiac cycle period). Data from two AI-VFM implementations, using PINNs trained with *NDop* = 200 and *NDop* = 15, are shown. As indicated by the dials symbolizing the cardiac cycle, the *NDop* = 200 AI-VFM is shown at frames (*•*) within the training set (*◦*). In contrast, the *NDop* = 15 AI-VFM is shown at frames outside of the training set. *A*) Radial velocity *v_r_*; *B*) Azimuthal velocity *v_θ_*; *C*) Flow vectors and color map of velocity magnitude; *D*) Flow vectors and color map of fluctuating pressure *p^′^*. Units for velocity and pressure are *cm/s* and *mmHg*, respectively.

The CFD-generated data, used as ground truth for evaluating AI-VFM, showed LV flow features in good agreement with previous simulations [56, 57] and clinical measurements [1]. As in the human LV, the CFD-simulated flows exhibited strong filling jets flanked by vortex rings and an emptying jet with prograde swirl. These patterns are the hallmarks of systolic and diastolic LV flow. Thus, although idealized, our CFD model provided an effective, physiologically representative benchmark flow for validating AI-VFM.

At time points with color-Doppler training data, AI-VFM inferred v*_r_* almost exactly (Figure 4A), which was expected since the *L_data_* loss directly penalized differences between the original and the reconstructed v*_r_* component. Moreover, outside of the timepoints with color-Doppler training data, AI-VFM also provided a faithful estimation of v*_r_*. The cross-beam velocities from AI-VFM closely matched the ground-truth CFD maps (Figure 4B). Because AI-VFM did not use any v*_θ_* data for training, this flow component estimation was not as precise as v*_r_*. Nevertheless, AI-VFM still recovered the dominant patterns of v*_θ_* even if some fine-scale features were not captured. The availability of color-Doppler training data at the interrogated temporal frame did not significantly influence the reconstruction of v*_θ_*. This result indicates that AI-VFM learned the dynamics of the flow to incorporate data across different time points despite the low temporal resolution in the training data.

An interesting feature of AI-VFM is that, by incorporating momentum equations in its loss functions, it can also infer flow pressure fields, p(r, θ, t). Pressure in incompressible flows is undetermined by a constant (i.e., pressure enters the governing equations 2 under a spatial gradient). Thus, to compare the inferred pressure maps with ground-truth maps from CFD (Figure 4D), we examined the pressure fluctuations p*^′^* = p *−* p by subtracting each snapshot’s mean pressure, p(t) = ^J^ p(r, θ, t)dΩ. The p*^′^*maps produced by AI-VFM agreed well with the ground-truth fields. During early filling, AI-VFM recovered the pressure fluctuations caused by the vortex ring flanking the mitral jet, and the vortex ring from the preceding beat as it impinged the LV apex. During late filling, AI-VFM reproduced pressure fluctuations induced by vortices as well as the adverse pressure gradient that decelerates the LV filling jet during this phase. Likewise, the pressure inferred by AI-VFM during ejection agreed with the CFD data, showing a favorable pressure gradient that accelerates the flow toward the aortic orifice.

The only noticeable difference between the pressure maps in AI-VFM and CFD was that AI-VFM produced smoother results, reaching lower p*^′^* peak values than CFD. This effect was more pronounced when AI-VFM was interrogated at temporal frames with no color-Doppler training data. PINNs are known to implicitly introduce smoothing by prioritizing learning the low-frequency features in the training data, a phenomenon known as spectral bias [58, 59]. To investigate whether this effect could be palliated by reducing the importance of the viscous terms in the PINN, we ran *dynamic* AI-VFM setting Re = *∞* in the Navier-Stokes loss terms (i.e., J_2_ and J_3_). The resulting flow maps were almost indistinguishable from those obtained using the Reynolds number value used in the CFD, Re = 4, 550 (Figure SI3).

#### 3.1.1 Point-wise Comparison of Velocity and Pressure Values

Scatter plots comparing v*_r_*, v*_θ_*, and p*^′^* from AI-VFM (y coordinate) and CFD (x coordinate) were built by pooling all the spatial points within the interrogated temporal frames, as indicated below. We overlaid contours of the joint probability density functions (pdfs) of VFM and CFD values and computed their r^2^ values. For reference, the outermost contour of these pdfs was chosen so that it left out 1% of the sampled points.

We tested the *dynamic* and *kinematic* versions of AI-VFM, along with the vanilla VFM method of Garcia *et al* [16], using all the CFD validation frames as input frames (e.g., N*_Dop_* = N*_BC_* = N*_cons_* = 200 for the PINNs). The results are displayed in the first row of Figure 5. The v*_r_* and v*_θ_* scatter plots from *kinematic* AI-VFM and vanilla VFM are shown as insets in the corners of each panel (top left and bottom right corners of Figures 5A-B, respectively). Consistent with the results shown in Figure 4, the scatter plots for v*_r_* aligned tightly along the identity line y = x for the three methods considered (Figure 5A), with r^2^ hovering around 0.99 in all the cases. Although the scatter plots of v*_θ_* experienced more dispersion than those of v*_r_*, *dynamic* AI-VFM still showed favorable agreement with the CFD ground-truth data, leading to r^2^ = 0.54. In comparison *kinematc* AI-VFM and vanilla VFM reached r^2^ = 0.31 and r^2^ = 0.25 respectively. Correspondence with ground-truth data was better for large v*_θ_* values, while more significant differences were observed around v*_θ_* = 0. Of note, *kinematic* AI-VFM and vanilla VFM showed more significant errors than *dynamic* AI-VFM when reconstructing negative values of v*_θ_*, suggesting that the Navier-Stokes constraint increased the accuracy of vector flow mapping. Pressure fluctuations from *dynamic* AI-VFM showed good agreement with CFD data (5*C*), with an overall r^2^ = 0.92. The agreement was particularly good for low and intermediate *|*p*^′^|* values, whereas AI-VFM tended to underestimate extreme values, consistent with the visualizations of Figure 4D.

**Figure 5:**
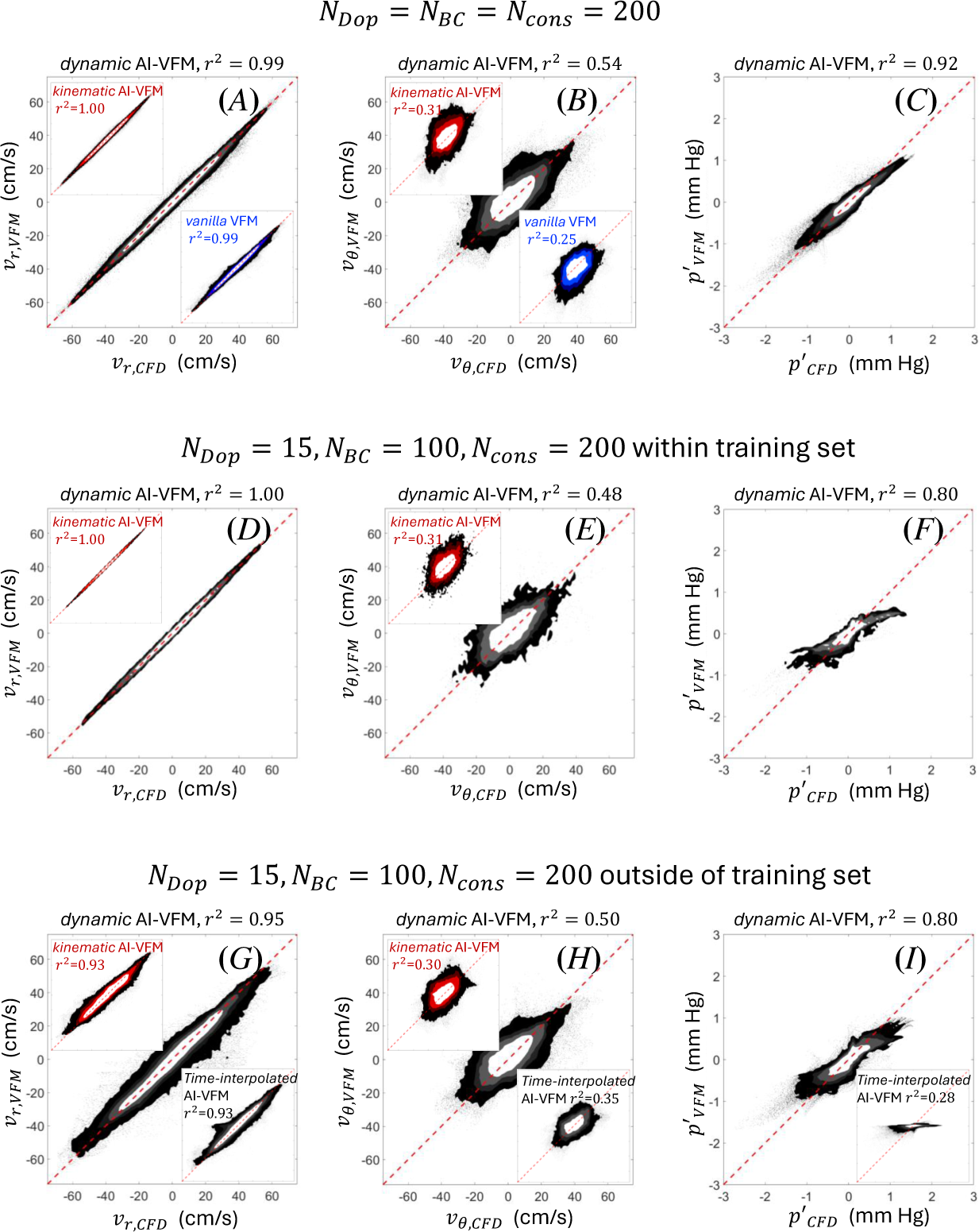
Point-wise comparison of velocity and fluctuating pressure values inferred by AI-VFM with ground-truth CFD data. Each panel shows scatter plots where the *x* and *y* coordinates are, respectively, the ground-truth CFD and AI-VFM values of flow variables sampled at uniformly spaced points inside the LV domain. The contour maps of joint probability densities of CFD and AI-VFM values are superimposed on each scatter plot. Units for velocity and pressure are *cm/s* and *mmHg*, respectively. *A-C*) AI-VFMs vs. CFD, *NDop* = *NBC* = *Ncons* = 200; *D-F*) AI-VFMs vs. CFD, *NDop*15*, NBC* = 100*, Ncons* = 200 within training set; *G-I*) AI-VFMs vs. CFD, *NDop* = 15*, NBC* = 100*, Ncons* = 200 outside of training set.

Next, we tested the ability of AI-VFM to learn hidden information about flow physics and recover velocity and pressure fields with finer temporal resolution than the training data. Thus, we ran AI-VFM using only N*_Dop_* = 15 frames with training v*_r_* data, but we imposed the boundary conditions and governing equations on more frames along the cardiac cycle (N*_BC_* = 100, N*_cons_* = 200). We then compared AI-VFM with ground-truth CFD data both at the N*_Dop_* = 15 within the training set (Figure 5D-F) and at the 185 frames outside of the training set (Figure 5G-I). Overall, the accuracy of *dynamic* AI-VFM decreased modestly when N*_Dop_* was lowered from 200 to 15, with r^2^ values for v*_r_*, v*_θ_* and p*^′^* varying from 0.99, 0.54, and 0.92 to 0.95-1.00, 0.48-0.50, and 0.80, respectively. The pressure fields were more sensitive to lowering N*_Dop_* than the velocity fields, and the shape of the pdfs suggests that peak pressure values were underestimated more significantly for N*_Dop_* = 15 than for N*_Dop_* = 200. Remarkably, *dynamic* AI-VFM had similar accuracy when interrogated within the training set (Figure 5D-F) and outside of it (Figure 5G-I).

*Kinematic* AI-VFM also conserved its performance when trained with N*_Dop_* = 15 instead of N*_Dop_* = 200; its accuracy was essentially unchanged both within the training range (top left corner insets of Figures 5D, *E*) and outside of it (top left corner insets of Figures 5G, *H*), except for a modest drop in r^2^ for v*_r_* from 1.00 to 0.93.

Finally, we ran *dynamic* AI-VFM using N*_out_* = N*_cons_* = N*_BC_* = N*_Dop_* = 15, and linearly interpolated its results in time at the 185 outside of the training dataset. This run aimed to evaluate the level of hidden flow information recovered by the PINN for different values of N*_BC_* and N*_cons_*. Their scatter plots are shown in the lower right corner insets of Figures 5G-I. The accuracy of the linearly interpolated AI-VFM maps dropped significantly, reaching values very similar to those obtained by *kinematic* AI-VFM. Overall, these results suggest that enforcing momentum balance at temporal frames without training data allows the PINN to learn hidden information about the flow dynamics, improving the accuracy of AI-VFM and providing temporal super-resolution. On the other hand, informing AI-VFM about flow kinematics (e.g., mass conservation) was insufficient to achieve super-resolution.

#### 3.1.2 Sensitivity Analyses

To assess the robustness of AI-VFM, we repeated the CFD-based validation for different spatial and temporal resolutions and transducer locations, as outlined in §2.8. To evaluate the distinct contributions of the Navier-Stokes and mass conservation constraints, we performed sensitivity analyses of the *dynamic* and *kinematic* implementations of AI-VFM. Figure 6 summarizes the results of these analyses, reporting NRMSEs of v*_r_*, v*_θ_*, and p*^′^* for *dynamic* AI-VFM, and NRMSEs of v*_r_* and v*_θ_* for *kinematic* AI-VFM.

**Figure 6:**
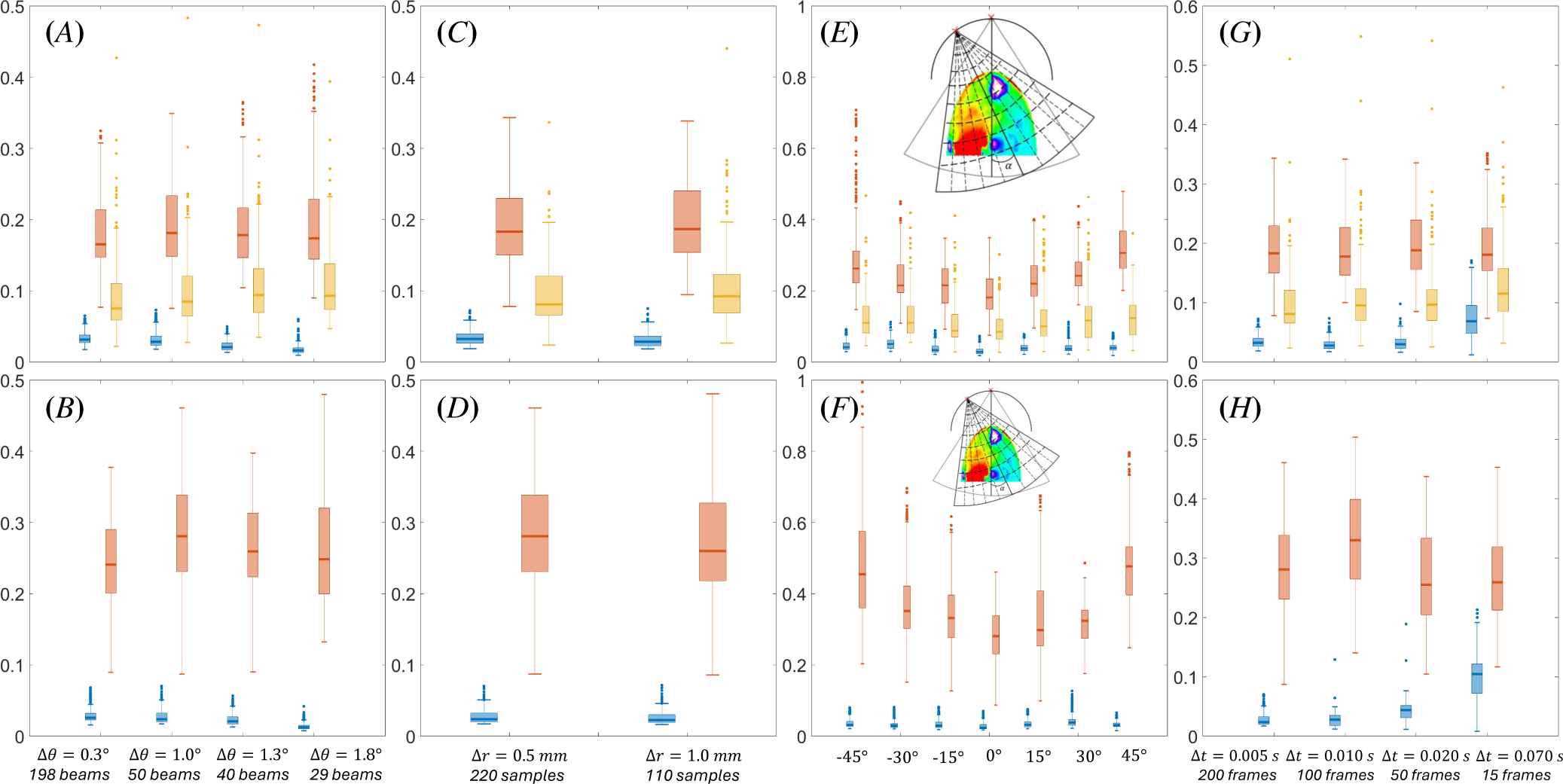
Normalized root mean squared error (NRMSE) of the radial velocity *v_r_* (blue), azimuthal velocity *v_θ_* (red), and pressure fluctuation *p^′^* (yellow) maps recovered by AI-VFM using CFD analysis as ground-truth data. The top row (panels *A*, *C*, *E*, and *G*) corresponds to *dynamic* AI-VFM whereas the bottom row (panels *B*, *D*, *F*, and *H*) corresponds to *kinematic* AI-VFM. *A*-*B*) NRMSE vs. spatial resolution in the azimuthal direction (i.e., angular step size Δ*θ*); *C*-*D*) NRMSE vs. spatial resolution in the radial direction (i.e., radial step size Δ*r*); *E*-*F*) NRMSE vs. ultrasound sector orientation (i.e., angle *α* between sector axis and LV long axis); *G*-*H*) NRMSE vs. temporal resolution of color-Doppler training sequence (i.e., *NDop* or Δ*t* = 1*/NDop*. When varying one parameter, all other parameters are kept fixed at nominal values (i.e., Δ*r* = 0.5*mm*, Δ*θ* = 1.0°, *N_dop_* = 200, and *α* = 0).

First, we varied the angular resolution while keeping nominal values for transducer positioning and radial and temporal resolutions. Since a standard color-Doppler acquisition has an angular step size Δθ *≈* 1.0°, we considered step sizes Δθ *≈* 0.3, 1.0, 1.3, and 1.8° in our sensitivity analysis. The NRMSEs of the reconstructed flow variables varied little with Δθ in the studied range for both *dynamic* AI-VFM (Figure 6A) and *kinematic* AI-VFM (Figure 6B). However, *dynamic* AI-VFM produced lower errors consistent with the results reported in the previous section. We also addressed the effect of radial resolution, varying the standard echocardiographic value, Δr *≈* 0.5mm, to Δr *≈* 1mm. Similar to the case of Δθ, this perturbation did not significantly modify the NRSME of the reconstructed flow variables in any of the two AI-VFM models (Figure 6C-D). Once again, *dynamic* AI-VFM was more accurate than *kinematic* AI-VFM over the range of studied resolutions.

We also examined the effect of transducer positioning by varying the angle α formed by the ultrasound sector axis and the CFD-LV long axis. For *dynamic* AI-VFM (Figure 6E), the NRMSEs of v*_r_* and p*^′^* were almost independent of α in the range [*−*45*^◦^*, 45*^◦^*] while the NRMSE of v*_θ_* experienced a shallow increase with *|*α*|*. In the case of *kinematic* AI-VFM (Figure 6F), the NRMSEs of v*_r_* were also insensitive to α. However, the reconstruction error of v*_θ_* increased more sharply with *|*α*|* in *kinematic* AI-VFM than in *dynamic* AI-VFM. In particular, the NRMSEs of *dynamic* AI-VFM at α = *±*45*^◦^* (0.26 and 0.31) were comparable to that of *kinematic* AI-VFM at α = 0 (0.28).

The temporal resolution of color-Doppler acquisitions is particularly sensitive to the width of the imaged sector. Because VFM works best when the color-Doppler sector encompasses the whole LV cavity, frame rates of 20 Hz or lower are not unusual in patients with dilated LVs. To address this issue, we studied AI-VFM performance while varying N*_Dop_* between 15 and 200. We found that *dynamic* AI-VFM was relatively insensitive to N*_Dop_* even if a modest increase in the NRMSEs of v*_r_* and p*^′^* was observed between N*_Dop_* = 50 and N*_Dop_* = 15 (Figure 6G). In contrast, *kinematic* AI-VFM was more sensitive to the temporal resolution of v*_r_*, experiencing an appreciable drop in the accuracy of its v*_θ_* reconstruction when N*_Dop_* decreased below 200, together with a more gradual deterioration of its v*_r_* reconstruction (Figure 6H).

### 3.2 Application to Clinical Acquisitions

This section demonstrates the feasibility of performing AI-VFM on clinical echocardiographic acquisitions. We illustrate this technique’s ability to correct aliasing artifacts in the color-Doppler data and reconstruct 2D maps of flow velocity vectors and pressure fluctuations. We also demonstrate that AI-VFM can restore missing data in clinical acquisitions, including entire color-Doppler frames. Since the validation and sensitivity analyses reported in previous sections suggest *dynamic* AI-VFM is more accurate and robust than *kinematic* AI-VFM, our clinical demonstration of AI-VFM focused on the *dynamic* implementation. Unless otherwise stated, we ran the PINNs using patient-specific values for N*_Dop_* and N*_BC_* in **Table** 1 given by the temporal resolution of each echocardiographic acquisition. In contrast, N*_cons_* = 200 frames were used to enforce mass conservation and momentum balance.

Phase wrapping artifacts are common in clinical color-Doppler images and can be particularly problematic during early diastole. During this phase, the flow velocity reaches large negative values inside the filling jet. If these velocity values are aliased, a large region of positive Doppler velocity, including the LV diastolic vortices, can be created, as shown in Figure

7*A*. This pattern can confound physics-unaware phase unwrapping algorithms, leading to jagged velocity contours that require artificial smoothing (Figure 7B) or, in worst-case scenarios, to the partial or total obliteration of the vortex signature in the color-Doppler signal (not shown). Figure 7C shows a clinical example where AI-VFM removed aliasing in the color-Doppler v*_r_* map, producing a smooth velocity field.

**Figure 7:**
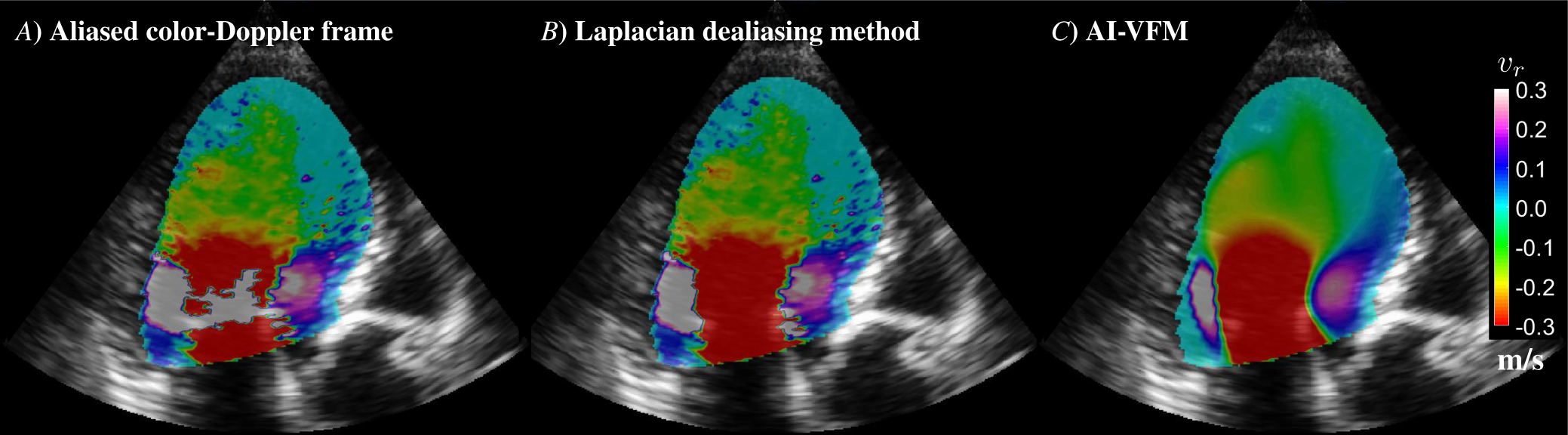
Color-Doppler dealiasing of radial velocity *vr* example in a clinical acquisition. *A*) Color-Doppler frame acquired during early diastole exhibiting an aliased region (white) in the central part of the inflow jet (red) that is merged with the positive velocity values associated with diastolic vortices. *B*) Color-Doppler frame dealiased using the one-step Laplacian method of Loecher *et al.*’s [55]. *C*) Color-Doppler output frame from AI-VFM. Velocity units are [m/s].

Figure 8 displays the velocity and pressure maps obtained by AI-VFM for the same patient considered in the previous figure. These maps are represented during early filling, late filling, and ejection. For completion, the raw color-Doppler maps at those time instants, some of which contain aliasing artifacts, are also represented (Figure 8A-C). Similar plots for two additional patients are shown in Supplementary Figures SI1 and SI2. The velocity vectors captured the early filling jet and its associated vortex ring (Figure 8D), which evolves into a single prograde swirling structure that occupies most of the LV chamber by late diastole (Figure 8E) and persists through ejection (Figure 8F), routing incoming blood toward the outflow tract. This spatiotemporal pattern is well-established as the representative LV flow pattern in the apical long-axis view [1]. In addition to the velocity field, AI-VFM calculated the pressure fluctuations inside the LV from clinical color-Doppler acquisitions (Figures

**Figure 8:**
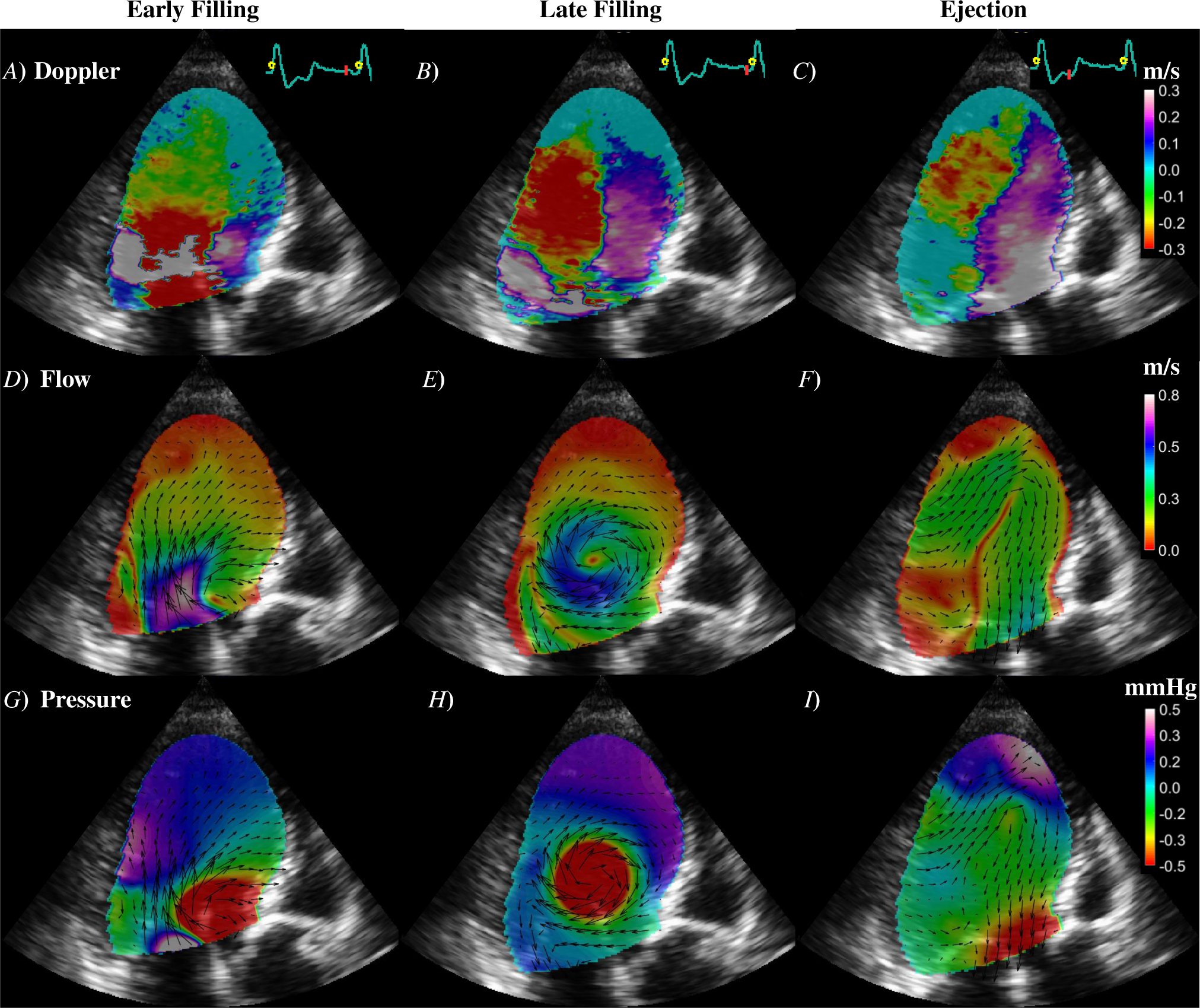
Example of clinical application of AI-VFM in patient #1. The first (panels *A*, *D*, and *G*), second (panels *B*, *E*, and *H*), and third (panels *C*, *F*, and *I*) columns display respectively early filling, late filling, and ejection. *A*-*C*) Raw color-Doppler training frames at indicated time points. *D*-*F*) Flow vectors and color map of velocity magnitude from AI-VFM at indicated time points. *G*-*I*) Flow vectors and pressure fluctuation at indicated time points. Velocity units are [*m/s*] and pressure units are [*mmHg*]. *NDop* = 17, *NBC* = 64, *Ncons* = 200 for *dynamic* AI-VFM.

8*G*-*I*). These pressure maps identified vortex cores since those are associated with local pressure minima, and agreed well with previously reported maps by secondary analyses of VFM maps [25].

To illustrate the capacity of AI-VFM to handle gappy images and regenerate the missing information, we re-ran the algorithm on the same patient, i.e., patient #1, as in Figure 8. However, this time, we created a large hole at the center of the color-Doppler sector in all the frames (N*_Dop_*) of the input sequence (Figure 9A-C). Despite the presence of these holes, the flow and pressure maps reconstructed by AI-VFM, shown in Figures 9D-I, were almost indistinguishable from those obtained when running AI-VFM on intact input images (Figures 8D-I), yielding r^2^ = 0.93, 0.87, 0.95 for v*_r_*, v*_θ_*, and p*^′^*, respectively N*_cons_* = 200 frames.

**Figure 9:**
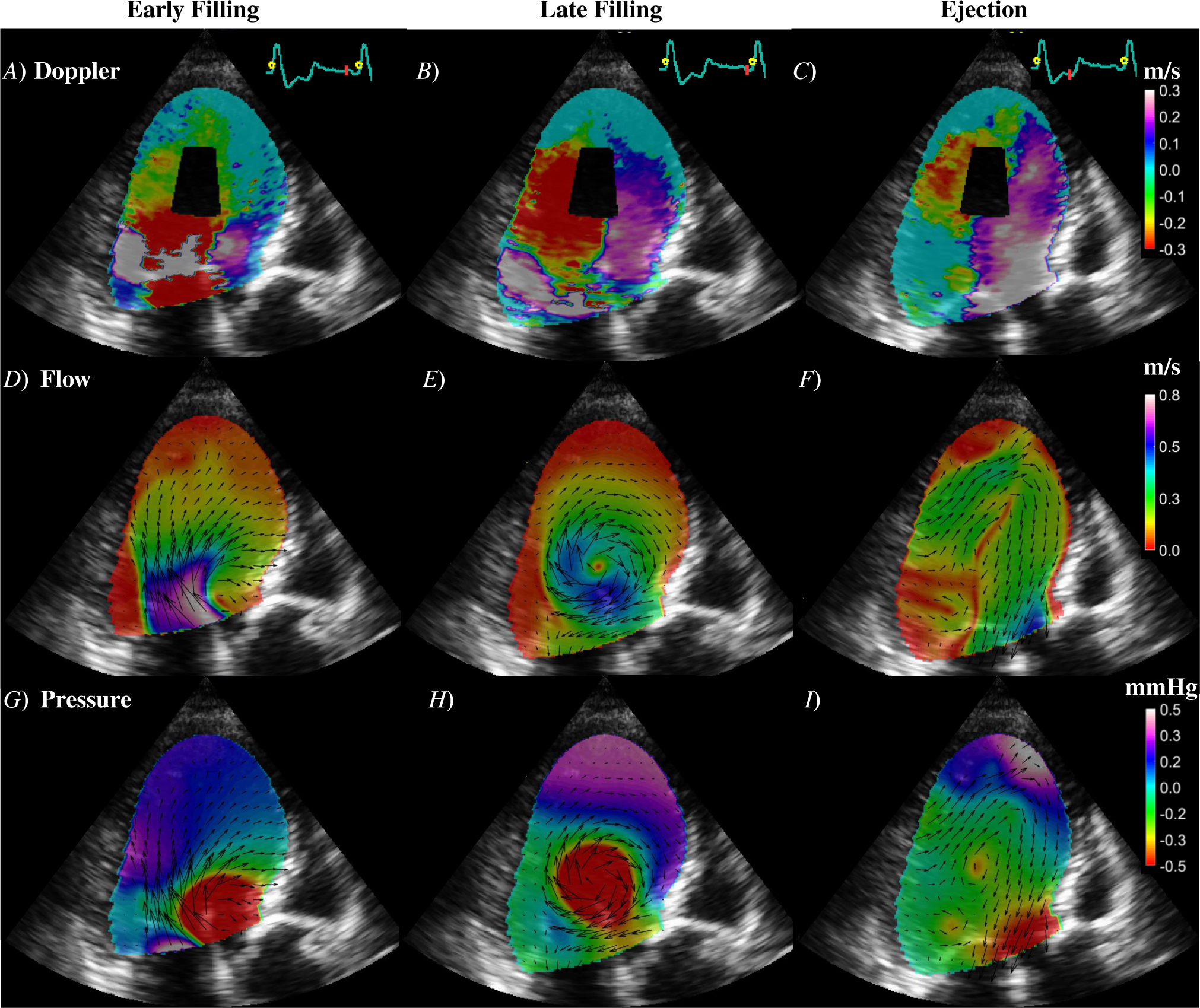
Example of clinical application of AI-VFM in patient #1 to color-Doppler sequences with holes inside the LV domain. The first (panels *A*, *D*, and *G*), second (panels *B*, *E*, and *H*), and third (panels *C*, *F*, and *I*) columns display respectively early filling, late filling, and ejection. *A*-*C*) Raw color-Doppler training frames at indicated time points displaying a large hole near the center of the LV chamber. *D*-*F*) Flow vectors and color map of velocity magnitude from AI-VFM at indicated time points. Note that AI-VFM reconstructs the velocity field inside the holes. *G*-*I*) Flow vectors and pressure fluctuation at indicated time points. Note that AI-VFM reconstructs the velocity field inside the holes. Velocity units are [*m/s*] and pressure units are [*mmHg*]. *NDop* = 17, *NBC* = 64, *Ncons* = 200 for *dynamic* AI-VFM.

Finally, we followed a leave-one-out approach to show that AI-VFM can also recover complete missing frames in clinical acquisition sequences. We created decimated versions of patient #1’s color-Dopppler acquisition by sequentially removing one of its N*_Dop_* color-Doppler frames. Then, we trained the PINN in *dynamic* AI-VFM using the remaining N*_Dop_ −* 1 frames, and used the color-Doppler frame excluded from training as ground-truth data for validation. Figure 10 compares ground-truth frames with the corresponding AI-VFM-reconstructed frames during early diastole, late diastole, and ejection, showing favorable visual agreement, and yielding r^2^ = 0.81 with respect to training with all available (N*_Dop_*) frames.

**Figure 10:**
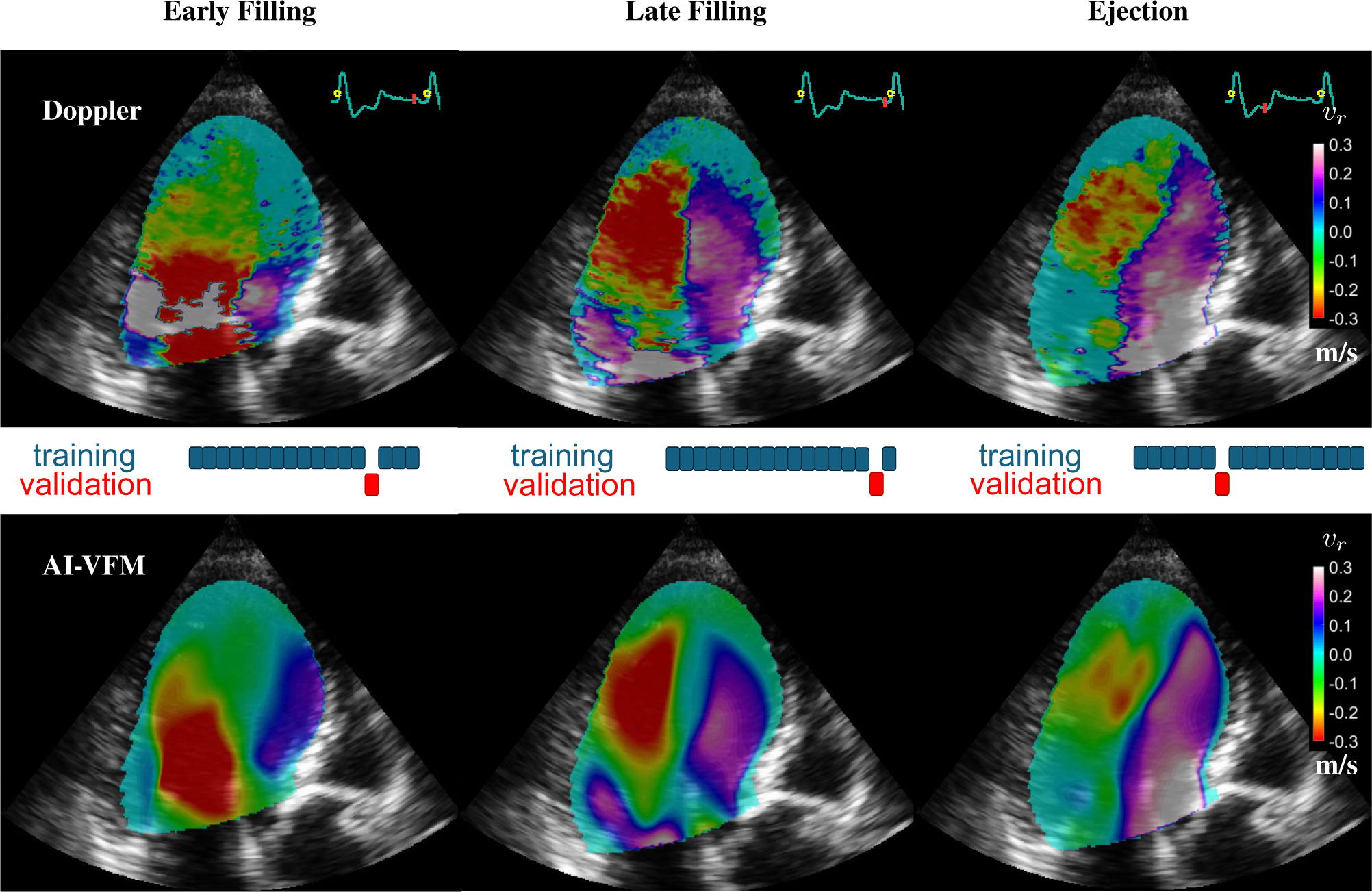
Leave-one-out validation of AI-VFM on the clinical color-Doppler sequence of patient #1. The three columns show results from three runs, where *dynamic* AI-VFM was trained on *NDop −* 1 color-Doppler frames, and one frame was dropped. The top image in each column is the dropped color-Doppler frame. The bottom image is the *vr* map recovered by AI-VFM for the instant of time corresponding to the dropped frame. Velocity units are [*m/s*]. *NDop* = 17, *NBC* = 64, *Ncons* = 200.

## 4 Discussion

Machine learning (ML) and AI are shaking the landscape of fluid mechanics research, offering unprecedented ability to study complex flows. In cardiovascular research, ML models are powerful tools to discover hidden relationships between clinical variables and diseases, offering valuable metrics for disease risk stratification [60–62]. The impact of deep learning (DL) models has been particularly significant [63]. Among these models, deep convolutional networks (CNNs) have proven effective at pattern recognition and feature extraction in complex flow data. Applied to cardiovascular flows, CNNs have proven helpful in resolving Doppler images aliasing artifacts [30], aortic wall shear stress estimation from 4D flow MRI [32], and real-time estimation of thrombotic risk in left atrial appendage [64]. Among the growing body of DL models, PINNs are well suited to study flow phenomena since they effectively incorporate the laws of physics as constraints in the learning process [34, 65]. PINNs have been used to enhance flow measurements from laboratory experiments [66] and medical imaging acquisitions [31, 36]. However, its applications to recover and visualize intracardiac flows from clinical data are scarce, and only preliminary approaches to the cardiovascular system have been recently conducted [31,67,68]. Some of these applications aim to use PINNs for improving cardiac simulations or modeling [29, 69, 70].

The application of DL models to reconstruct LV flow from color-Doppler data is just beginning to be explored but offers significant promise of improvement over traditional VFM methods [35, 71]. This study centered around PINNs as the means of flow reconstruction, focusing on the performance gains obtained by incorporating physical constraints related to flow kinematics (i.e., mass conservation) and dynamics (i.e., momentum balance). Unprecedented in VFM, we included momentum balance in the AI-VFM PINNs via loss terms that measure the residuals of the planar Navier-Stokes equations. Since these equations include pressure gradients, the pressure was an output variable of the PINNs, allowing AI-VFM to recover pressure fluctuations inside the LV. Furthermore, informing AI-VFM with flow dynamics allowed this modality to provide temporal super-resolution and recover entire missing frames from an acquisition sequence.

Our comparative analyses of *kinematics*-informed vs. *dynamics*-informed AI-VFM suggested that enforcing momentum balance in the PINN increased the accuracy and robustness of VFM. Overall, *dynamic* AI-VFM was more accurate than *kinematic* AI-VFM, which in turn was more accurate than the vanilla VFM implementation that integrates the continuity equation separately along each arc of the Doppler sector [16]. Moreover, our sensitivity analyses showed *dynamic* AI-VFM to be less sensitive to the input data’s spatial and temporal resolutions and the ultrasound sector’s orientation. The sensitivity to ultrasound sector alignment is particularly relevant since VFM relies on the inflow and outflow jets of the LV being captured by the color-Doppler velocity component. Because those jets are primarily aligned with the LV long axis, the performance of VFM deteriorates as the angle α between the LV long axis and the ultrasound sector increases [16]. Still, we found *dynamic* AI-VFM to perform as accurately for α = 45° as *kinematic* AI-VFM for α = 0.

The Navier-Stokes equations include acceleration and viscous terms representing physical mechanisms distinct from mass conservation. Informing AI-VFM with flow acceleration allowed this method to connect information from different temporal frames within each training set. Consequently, *dynamic* AI-VFM produced temporally super-resolved outputs with velocity and pressure maps at time points without color-Doppler training data. When forcing *dynamic* AI-VFM to linearly interpolate between training frames without enforcing momentum balance inside the intervals defined by these frames, its performance degenerated to that of *kinematic* AI-VFM. This experiment supports the idea that enforcing momentum balance is crucial to achieving temporal super-resolution. From a practical point of view, handling color-Doppler input sequences with few more than a dozen frames spanning one single heartbeat facilitates the clinical adoption of VFM because it reduces the time patients need to hold their breath while being scanned from *∼* 10 s to *∼* 1 s. In our experience, breath-holding has been a limiting factor of VFM, especially in patients with severely impaired cardiac function. This feature also opens the door to applying AI-VFM to patients with heart rhythm disorders.

In color-Doppler acquisitions, the Nyquist velocity limit beyond which the Doppler phase wraps around 2π, is dictated by the ultrasound pulse repetition frequency (PRF). Increasing the PRF allows for detecting higher flow velocities but compromises the scanning depth of the acquisition. Consequently, color-Doppler sequences for VFM are often acquired using Nyquist velocities lower than the peak velocities found in the inflow and outflow jets of the LV. Thus, these sequences cancontain significant aliasing artifacts. Since aliasing artifacts are common to other flow imaging modalities like phase-contrast MRI, they have received significant attention, and standalone algorithms have been developed for their removal [30, 55, 72, 73]. In a nutshell, these algorithms use knowledge about phase-wrapping, training data, and/or smoothness constraints to decide whether jumps in velocity between neighboring pixels are caused or not by phase wrapping. Using physical constraints should help make this decision but had not been attempted in color-Doppler echocardiography so far. In AI-VFM, we followed an approach recently proposed for 4D-flow MRI [36]. We made the data loss term in the PINN insensitive to 2π jumps in the phase of the color-Doppler signal and let the physics constraints in the PINN ensure the regularity of the output flow maps. We found this approach was sufficient to correct aliasing artifacts in several clinical color-Doppler acquisitions without introducing any explicit dealiasing step in the algorithm. We also found that the dealiased color-Doppler maps generated by AI-VFM compared well with those obtained with standalone methods informed by phase wrapping rules [55].

In addition to aliasing, clinical color-Doppler images sometimes include sizeable areas of unreliable data caused by insufficient signal power, device-related ultrasound reflections [74], etc. These artifacts are more challenging to remedy than aliasing because they are not governed by a simple phase-wrapping rule. We have shown that AI-VFM seamlessly tolerates large holes in the input data, using the hidden information about flow physics to restore the flow inside these holes. Therefore, AI-VFM offers a new method to analyze artifact-ridden images as long as the unreliable pixels in each image can be labeled and excluded from training. Future efforts should focus on the automation of unreliable pixel labeling and the design of data loss terms that tolerate such pixels, similar to the 2π-insensitive data loss term used in this work.

The inclusion of viscous terms in the PINN loss should contribute to regularizing the velocity fields in ways consistent with flow physics. The flexibility of the PINN framework allows for blood viscosity to be an output parameter [65], so no prior knowledge of this parameter is needed. Nevertheless, our results suggest that spectral bias in the PINNs [58, 59] created at least as much smoothing as the viscous terms for physiologically representative Reynolds number values. Therefore, the *effective* Re of the AI-VFM-reconstructed flow maps was lower than the actual Re of the flow and, consequently, the peak values of, e.g., pressure fluctuations, were underestimated by AI-VFM. On the other hand, except for the presence of imaging noise, the clinical color-Doppler maps do not exhibit the fine-scale flow patterns expected to be found in LV flow at physiologically representative Reynolds number values of Re *≈* 4, 000. This averaging effect in color-Doppler maps was previously noted by Seo *et al* [56] by comparing with their CFD simulations. Therefore, while further efforts are warranted to mitigate spectral bias in AI-VFM [59, 75], the clinical application of this technique should not be hindered by this limitation. Furthermore, our leave-one-out validation of AI-VFM on clinical acquisitions suggests that the PINN captured most if not all the flow features in the color-Doppler maps.

The PINN approach followed in AI-VFM ensures this technique is trained using data from one patient at a time, bypassing the need for training on large pre-existing data sets. PINNs are generally less sensitive to common issues of DL models like the misrepresentation of features underrepresented in the training set [26, 60, 61]. While the need for re-training the PINN for every single patient may seem computationally expensive, this demand is manageable in VFM given the relatively light weight of the input 2D image sequences. Currently, AI-VFM converges reasonably within 15 minutes on an off-the-shelf contemporary graphics processing unit (GPU), and its accuracy can be improved by an additional 10% by extending training to one hour. Moreover, we did not make an exhaustive effort to fine-tune the number of PINN layers/neurons, batch size, training iterations, and the weights of different loss terms, leaving room for further accelerations in convergence. Considering these factors, the rapid progress in GPU architecture, and the Tensorflow software engine used by the PINNs [76], we expect that further reductions in compute time that could bring AI-VFM closer to real-time execution are well within reach. Future efforts for accelerating AI-VFM and mitigating the *ad hoc* choices of weight coefficients will include the implementation of dynamic weights of the loss terms as proposed in the literature [59, 77, 78].

The fully-connected neural network configuration in AI-VFM provides a flexible framework that can be expanded without great difficulty to include additional input and output variables, as well as physical constraints. In the field of echocardiography, this flexibility could be exploited to assimilate flow fields from blood speckle imaging [14, 15], a technique based on high-frequency ultrasound that measures 2D blood flow vectors but often produces gappy data. In addition, future directions will explore including additional physical constraints in the PINNs of AI-VFM to map LV blood residence time [79] or the concentration of coagulation species [80].

## 5 Conclusion

This manuscript describes AI-VFM, a new vector flow mapping (VFM) method enabled by recent advances in artificial intelligence (AI). AI-VFM uses physics-informed neural networks (PINNs) encoding mass conservation, momentum balance, and boundary conditions to recover intraventricular flow and pressure fields from standard echocardiographic scans. AI-VFM performs phase unwrapping and recovers missing data in the form of spatial and temporal gaps in the input color-Doppler data, thereby producing super-resolution flow maps. AI-VFM is solely informed by each patient’s flow physics; it does not utilize explicit smoothness constraints or incorporate data from other patients or flow models. AI-VFM shows good validation against ground-truth data from CFD, outperforming traditional VFM methods as well as similar PINN-based VFM formulations relying exclusively on mass conservation.

## Acknowledgments

This work was supported by the National Institutes of Health under grants 1R01HL160024 and 1R01HL158667, the National Science Foundation Graduate Research Fellowship under grant No.DGE-2140004, the Spanish Research Agency and the European Regional Development Fund under grant PID2019-107279RB-I00, the Comunidad de Madrid and the European Regional Development Fund under grant Y2018/BIO-4858 PREFI-CM, and the Instituto de Salud Carlos III under grants PI20/00587-AI4NHEM, PI21/00274-PACER, and DTS/1900063-ISBIFLOW.

## Supporting Information

Figures SI1 and SI2 in this section demonstrate the clinical application of AI-VFM to two additional patients (#2 and #3, see Table 1 in the main text). Figure SI3 compares CFD and AI-VFM data when the PINN is informed with and without viscous terms.

**Figure SI1:**
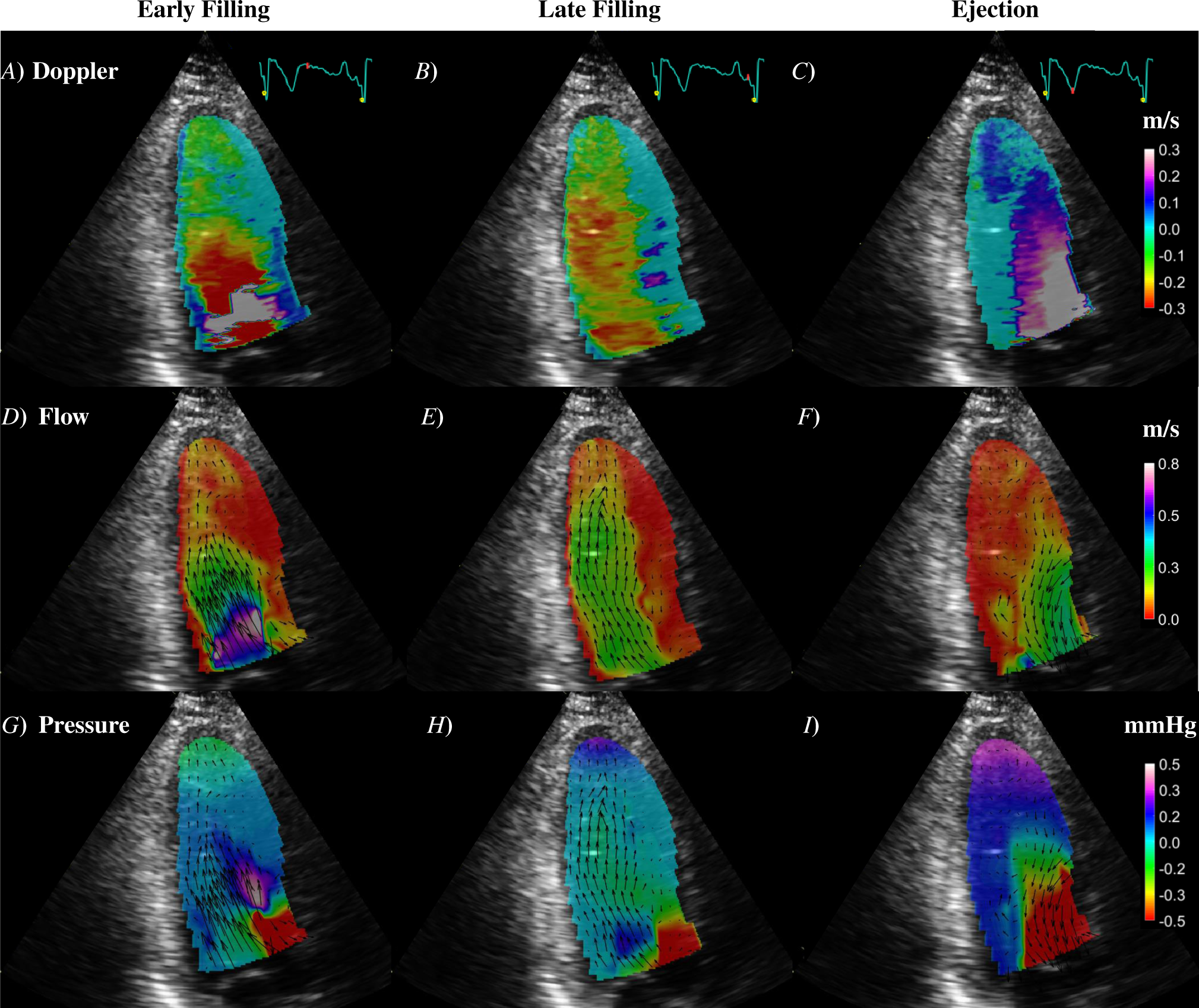
Example of clinical application of AI-VFM in patient #2. The first (panels *A*, *D*, and *G*), second (panels *B*, *E*, and *H*), and third (panels *C*, *F*, and *I*) columns display respectively early filling, late filling, and ejection. *A*-*C*) Raw color-Doppler training frames at indicated time points. *D*-*F*) Flow vectors and color map of velocity magnitude from AI-VFM at indicated time points. *G*-*I*) Flow vectors and pressure fluctuation at indicated time points. Velocity units are [*m/s*] and pressure units are [*mmHg*]. *NDop* = 19, *NBC* = 74, *Ncons* = 200 for *dynamic* AI-VFM..

**Figure SI2:**
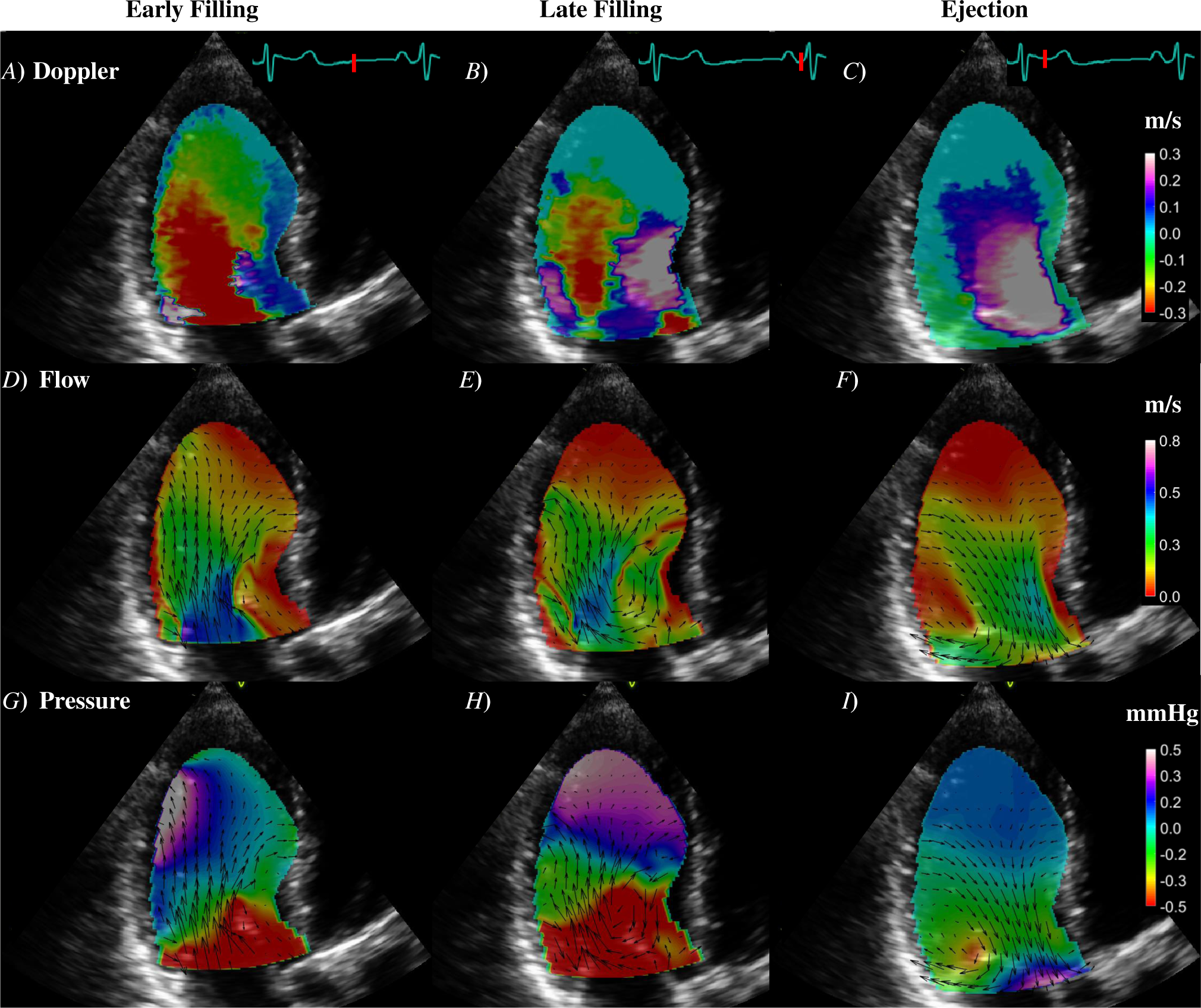
Example of clinical application of AI-VFM in patient #3. The first (panels *A*, *D*, and *G*), second (panels *B*, *E*, and *H*), and third (panels *C*, *F*, and *I*) columns display respectively early filling, late filling, and ejection. *A*-*C*) Raw color-Doppler training frames at indicated time points. *D*-*F*) Flow vectors and color map of velocity magnitude from AI-VFM at indicated time points. *G*-*I*) Flow vectors and pressure fluctuation at indicated time points. Velocity units are [*m/s*] and pressure units are [*mmHg*]. *NDop* = 19, *NBC* = 83, *Ncons* = 200 for *dynamic* AI-VFM..

**Figure SI3:**
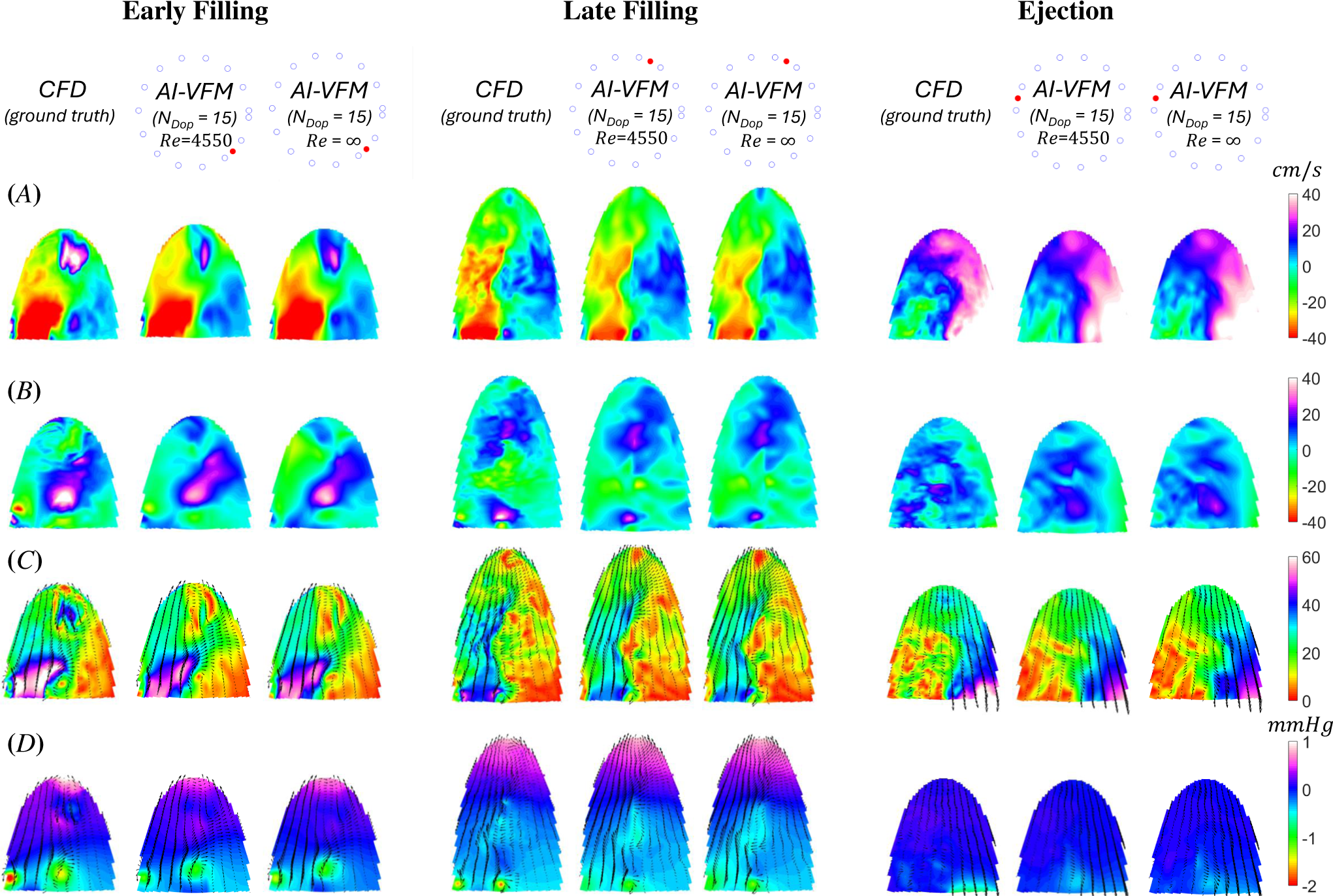
Velocity and pressure maps inferred by AI-VFM compared to ground-truth CFD data at early filling, late filling, and ejection (i.e., *t* = 0.85, 0.45, 0.21*T*, where *T* is the cardiac cycle period). Data from two AI-VFM implementations, using PINNs trained with *Re* = 4, 550 and *Re* = *∞*, are shown. Both are trained with *NDop* = 15, *NBC* = 100, and *Ncons* = 200. As indicated by the dials symbolizing the cardiac cycle, bot are shown at frames (*•*) outside of the training set (*◦*). *A*) Radial velocity *v_r_*; *B*) Azimuthal velocity *v_θ_*; *C*) Flow vectors and color map of velocity magnitude; *D*) Flow vectors and color map of fluctuating pressure *p^′^*. Units for velocity and pressure are *cm/s* and *mmHg*, respectively.

